# Ndufs4 inactivation in glutamatergic neurons reveals swallow-breathing discoordination in a mouse model of Leigh Syndrome

**DOI:** 10.1101/2024.09.11.612506

**Authors:** Alyssa Huff, Luiz Marcelo Oliveira, Marlusa Karlen-Amarante, Favour Ebiala, Jan Marino Ramirez, Franck Kalume

## Abstract

Swallowing, both nutritive and non-nutritive, is highly dysfunctional in children with Leigh Syndrome (LS) and contributes to the need for both gastrostomy and tracheostomy tube placement. Without these interventions aspiration of food, liquid, and mucus occur resulting in repeated bouts of respiratory infection. No study has investigated whether mouse models of LS, a neurometabolic disorder, exhibit dysfunctions in neuromuscular activity of swallow and breathing integration. We used a genetic mouse model of LS in which the NDUFS4 gene is knocked out (KO) specifically in Vglut2 or Gad2 neurons. We found increased variability of the swallow motor pattern, disruption in breathing regeneration post swallow, and water-induced apneas only in Vglut2 KO mice. These physiological changes likely contribute to weight loss and premature death seen in this mouse model. Following chronic hypoxia (CH) exposure, swallow motor pattern, breathing regeneration, weight, and life expectancy were not changed in the Vglut2-Ndufs4-KO CH mice compared to control, indicating a rescue of phenotypes. These findings show that like patients with LS, Ndufs4 mouse models of LS exhibit swallow impairments as well as swallow-breathing dyscoordination alongside the other phenotypic traits described in previous studies. Understanding this aspect of LS will open roads for the development of future more efficacious therapeutic intervention for this illness.

## Introduction

Swallowing serves two physiological functions: passing food to the stomach for nutrition, and pharyngeal clearance for airway protection. Difficulty feeding and swallowing are some of the first symptoms of Leigh Syndrome (LS) (1–3). LS is the most frequent form of pediatric mitochondrial disease, affecting 1 in every 40,000 births (4). Clinical manifestations of LS typically appear in childhood, within the first 2 years of life, though adult-onset of the disease has also been reported (5–9). These manifestations of the disease are marked by symmetrical lesions to the basal ganglia, brainstem, and spinal cord which are associated with multiple and heterogenous symptoms such as psychomotor regressions, hypotonia, dystonia, seizures, movement disorders, ataxia, feeding and swallowing disorders, respiratory disorders, neuropathy, and myopathy (10, 11).

It has been reported that up to 45% of LS patients have some type of feeding and swallowing disorder (2, 9, 12) described as “feeding difficulties” (10, 12), “difficulty or inability to suck” (2, 3), “dysphagia” (2, 6), “decreased pharyngeal reflex” (7), “swallowing dysfunction” (9), and “regurgitation or vomiting during or after feeding” (2). These impairments contribute to a high risk of developing respiratory infections (aspiration pneumonia), respiratory failure, a failure to thrive (6, 13), which are all common causes of death in LS (2, 4, 8, 9, 14). The lack of understanding in how mitochondrial dysfunction disrupts swallow-breathing neurocircuitry has resulted in a limited number of therapies designed to combat malnutrition and airway protection. Gastrostomy (G-tube) and tracheostomy placement are common procedures used to promote growth by increasing caloric and fluid intake, and to safely clear secretions, respectively (2, 5, 9, 13, 15, 16). Although these procedures have shown positive impacts on growth, nutrition, and respiratory health, they are associated with several complications, including local infection, vomiting and gastroesophageal reflux (17). In addition, it is worth noting that these and other therapeutic options to address feeding and swallow difficulties available to date are not curative, their goal is to reduce neurodegeneration, respiratory disturbances, and improve feeding using G-tube (5).

LS is associated with more than 80 genes of the nuclear or mitochondrial genome, making its diagnosis and treatment complicated (10). Mutation to the NADH dehydrogenase (ubiquinone reductase) iron-sulfur protein 4 (NDUFS4) gene causes early-onset severe LS that results in early death (18). NDUFS4 encodes for the NADH: ubiquinone oxioreductase subunit S4 of mitochondrial complex I, a key structural component for the assembly, stability and activity of complex I (19). Mice harboring a whole-body homozygous KO of this gene constitute a well validated model of the disease and has been shown to exhibit several key clinical phenotypic traits of LS (20). Interestingly, Bolea et al. recently demonstrated that different neuronal cell types drive distinct sets of LS symptoms. In particular, they showed that excitatory, not inhibitory, neurons drive the development of motor and breathing deficits in the LS Ndufs4 KO mice (11). These phenotypes were observed in mice with Ndufs4 KO restricted to Vglut2 neurons (Vglut2-Ndufs4-KO). These Vglut2-Ndufs4-KO mice also exhibited decreased body weight, food intake, and life expectancy (11), indicating a possible feeding and swallowing disorder likely leading to failure to thrive. Swallow and breathing are both dependent of motor function and are tightly coordinated to ensure proper gas exchange during breathing and prevent aspiration during swallowing (21). However, direct assessments of these functions have yet to be conducted in these LS mice.

Here we investigated and described swallow motor activity and its coordination with breathing in healthy and Vglut2-Ndufs4-KO mice. We hypothesize that an LS-causing mutation of the NDUFS4 gene can disrupt not only normal breathing but also swallow motor activity. These phenotypes may arise from the mutation related mitochondrial and excitability dysfunctions in excitatory neurons of swallow and breathing medullary regions such as the Ventral Respiratory Column (VRC), postinspiratory complex (PiCo), and the nucleus of the solitary tract (NTS) (22–24). Further, we hypothesized that exposure to chronic hypoxia, a therapeutic approach that has proven to protect against oxidative stress and neurodegeneration in LS models (25, 26), will prevent swallow and breathing abnormalities, rescue body weight and life span in Vglut2-Ndufs4-KO mice.

## Methods

### Animals

Both adult male (P68-236, average P142) and female (P64-237, average P144) mice were bred at Seattle Children’s Research Institute (SCRI) and used for all experiments. All data pertaining to sex specific measured parameters are in tables 4-6. Our healthy control mice consisted of C57BL/6J obtained from The Jackson Laboratory (Stock No:000664) and Ndufs4-Vglut2cre negative mice. The experimental mice were Ndufs4-Vglut2 cre positive and Gad2 cre positive (Vglut2cre-Ndufs4-KO and Gad2cre-Ndufs4-KO). Mice with conditional deletion of Ndufs4 in Vglut2-expressing glutamatergic neurons were generated by crossing mice with one Ndufs4 allele deleted and expressing a codon-improved Cre recombinase (iCre) under the Slc17a6 promoter (Slc17a6Cre, Ndufs4D/+) to mice with two floxed Ndufs4 alleles (Ndufs4lox/lox). Mice with conditional deletion of Ndufs4 in Gad2-expressing GABAergic neurons (experimental mice) were generated by crossing mice with one Ndufs4 allele deleted and expressing a codon-improved Cre recombinase (iCre) under the Gad2 promoter (Gad2Cre, Ndufs4D/+) to mice with two floxed Ndufs4 alleles (Ndufs4lox/lox). Ndufs4lox/lox were generated by Quintana et al., 2010; Kruse et al., 2008. Slc17a6Cre (BAC-Vglut2::Cre) (Borgius et al., 2010) mice were generated by Ole Kiehnor. Gad2Cre/+ (Gad2-IRES-Cre) (The Jackson Laboratory Stock No:028867). All mice were on a C57BL/6J background. Genotype of the offspring and absence of ectopic recombination (i.e. presence of recombination bands in tail DNA samples) was determined by PCR analysis. Offsprings were group housed with ad libitum access to food and water in a temperature controlled (22 ± 1°C) facility with a 12h light/dark cycle. All experiments and animal procedures were approved by the Seattle Children’s Research Institute’s Animal Care and Use Committee and were conducted in accordance with the National Institutes of Health guidelines.

### Chronic Hypoxia

Vglut2-Ndufs4-KO (average age P52) and control (average age P45) mice were kept in collective cages with food and water ad libitum placed inside plexiglass chambers filled with 11% O_2_ 24 hours a day on light cycle in a 12hr light/dark cycle. Ten of the 13 Vglut2-Ndufs4-KO mice were in CH for an average 180 days, while 3 mice had to be removed early due to development of an eye condition and were removed at day 114 and 64. Control mice stayed in CH for an average of 120 days.

### *In Vivo* Experiments

The same experimental protocol was performed for all control, Vglut2:Ndufs4KO and Gad2:Ndufs4KO. However, the age of each experimental group was different due to some physiological and experimental factors. Experimental protocol for physiological assessments requires alive, yet anesthetized mice, resulting in artificially picking their end date. For all control mice, the age at experiment was picked to match the age of its Vglut2cre-Ndufs4-KO counterpart. Gadcre-Ndufs4-KO mice age at experiment depended on symptoms and seizure onset. Criteria for Vglut2cre-Ndufs4-KO RA mice end date was onset of symptoms, little to no movement in the cage, cold to touch and/or a body condition score of 1. For Vglut2cre-Ndufs4-KO CH mice experiments were done following 6 months of CH exposure. Three of those mice had to be removed from CH and experiment performed, 1 at 4 months and 2 at 2 months, due to development of an eye condition. Adult mice were injected with Urethane (1.5 g/kg, i.p. Sigma Aldrich, St. Louis, MO, USA) and secured supine on a custom surgical table. Core temperature was maintained through a water heating system (PolyScience, Niles, IL, USA) built into the surgical table. Mice were then allowed to spontaneously breathe 100% O_2_ for the remainder of the surgery and experimental protocol. Adequate depth of anesthesia was determined via heart and breathing rate, as well as lack of toe pinch response every 15 minutes. A supplemental dose of 0.01mL of Urethane was given to maintain adequate anesthetic depth, when necessary.

Bipolar electromyograms (EMG) electrodes were placed in the costal diaphragm to monitor respiratory rate and heart rate throughout the experiment. The trachea was exposed through a midline incision and cannulated caudal to the larynx with a curved (180 degree) tracheal tube (PTFE 24 G, Component Supply, Sparta, TN, USA). The recurrent laryngeal nerve (RLN) was carefully dissected away from each side of the trachea before the cannula was tied in and sealed with super glue to ensure no damage to the RLN. A tube filled with 100% O_2_ was attached to the cannulated trachea to provide supplemental oxygen throughout the experiment. As previously published (Figure. 6a (22)), the XII and X nerves were isolated unilaterally, cut distally, and their activity was recorded from a fire-polished pulled borosilicate glass tip (B150-86-15, Sutter Instrument; Novato, CA, USA) filled with artificial cerebral spinal fluid (aCSF; in mM: 118 NaCl, 3 KCl, 25 NaHCO_3_, 1 NaH_2_PO_4_, 1 MgCl_2_, 1.5 CaCl_2_, 30 D-glucose) equilibrated with carbogen (95% O_2_, 5% CO_2_), connected to the monopolar suction electrode (A-M Systems, Sequim, WA, USA) and held in a 3D micromanipulator (Narishige, Tokyo, Japan). Multiple bipolar EMGs, using 0.002” and 0.003” coated stainless steel wires (A-M Systems, Sequim, WA, USA, part no.790600 and 791000 respectively), simultaneously recorded activity from several swallow and respiratory-related muscle sites. According to the techniques of Basmajian and Stecko (27), the electrodes were placed using hypodermic needles 30G (part no 305106, BD Precision Glide ™, Franklin Lakes, NJ, USA) in the *submental complex*, which consists of the geniohyoid, mylohyoid and anterior digastric muscles, to determine swallow activity. The *laryngeal complex,* consisting of the posterior cricoarytenoid, lateral, transverse and oblique arytenoid, cricothyroid and thyroarytenoid muscles, to determine laryngeal activity during swallow, as well as postinspiratory activity. The *costal diaphragm*, used to measure the multifunctional activity for both inspiration, *Schluckatmung*, a less common diaphragmatic activation during swallow activity, and airway protective behaviors such as aspiration reflex (AspR) (Figure 1B, figure supplement 1, Figure 2C). Swallow was stimulated by injecting 0.1cc of water into the mouth using a 1.0 cc syringe connected to a polyethylene tube. At the end of the experiment mice were euthanized by an overdose of anesthetic followed by trans-cardial perfusion (see histology section below).

**Figure 1.**
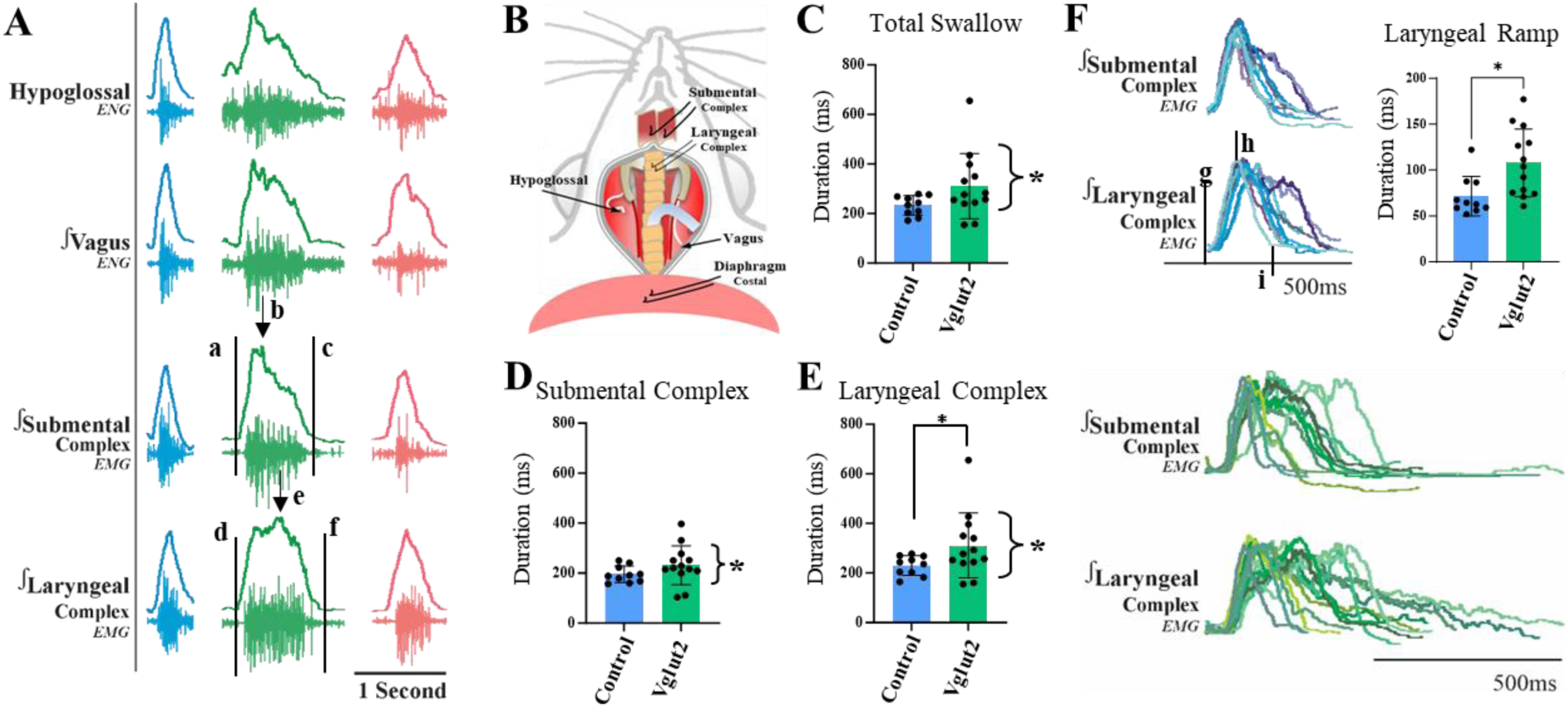
Alteration in, and variability of swallow motor pattern in Vglut2-Ndufs4-KO mice exposed to room air. A) Representative traces of water triggered swallow in control (blue), Vglut2-Ndufs4-KO (green), and Gad2-Ndufs4-KO (pink). Total swallow duration: f-a; submental complex (SC) duration: c-a; laryngeal complex (LC) duration: f-d; and swallow sequence: e-b. B) Graphical depiction of the preparation with location of all recorded muscles and nerves. The F test to compare variances indicated a statistically significant variance between Vglut2-Ndufs4-KO and control room air (RA) mice in C) the total swallow duration, D) SC duration, and E) LC duration indicated with the bracket and asterisk to the right of the bar graphs. F) Overlay of representative trace of a swallow from each animal in control RA (blue, top) and Vglut2-Ndufs4-KO RA (green, bottom) to illustrate the variable swallow motor patten in the Vglut2-Ndufs4-KO mice but not the control mice. Ramp duration was calculated as h-g in both SC and LC; and decay duration from i-h. LC in Vglut2-Ndufs4-KO mice have a significantly longer duration (E), likely due to the significantly longer ramping of the muscle activity (F) compared to the control RA mice.

**Figure 2.**
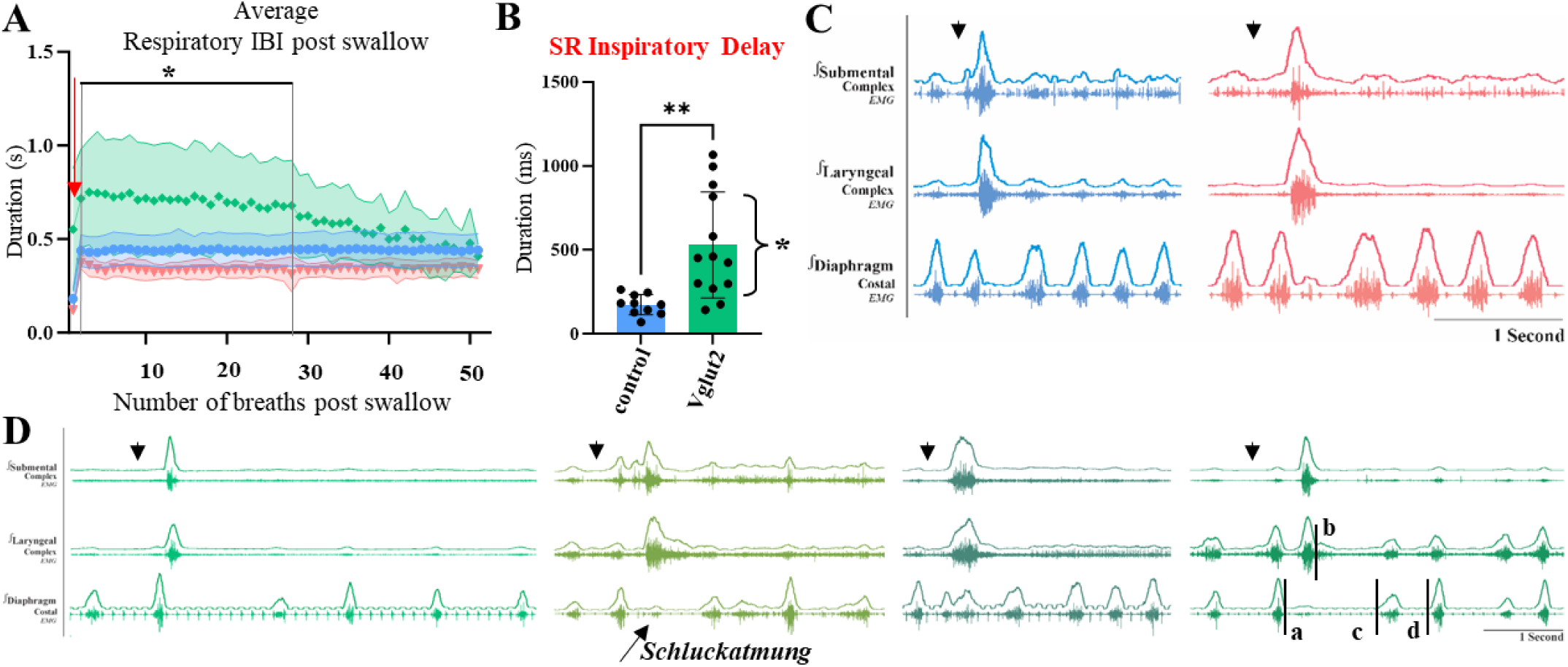
Disruption of respiratory rhythm regeneration following a swallow in Vglut2-Ndufs4-KO room air mice. Respiratory rhythm regeneration was determined by calculating the inter-burst-interval (IBI) for 50 cycles following a swallow. A) Plot of the average diaphragm IBI following a swallow for control (blue), Vglut2-Ndufs4-KO (green), and Gad2-Ndufs4-KO (pink) exposed to room air. Vglut2-Ndufs4-KO mice have significantly longer IBI compared to control mice from breaths 1-28 and increased variability (shaded area). The red arrow indicates the first breath post swallow termed B) swallow related (SR) inspiratory delay. Vglut2-Ndufs4-KO mice have a significantly longer and variable (asterisk on the side) duration in which breathing regenerates following a swallow. C) Representative traces of control (blue) and Gad2-Ndufs4-KO (pink) of breathing regeneration following a swallow induced by water (down arrow). D) Representative traces of 4 Vglut2-Ndufs4-KO mice indicating the variability of breathing regeneration and 4 cycles following a swallow. *Schluckatmung*, a less known behavior describes diaphragm activity during swallow (55, 56). SR expiration duration d-a, SR inspiratory delay c-b, diaphragm IBI d-c.

## Analysis

All electroneurogram (ENG) and EMG activity were amplified and band-pass filtered (0.03 – 1 KHz) by a differential AC Amplifier (A-M System model 1700, Sequim, WA, USA), acquired in an A/D converter (CED 1401; Cambridge Electronic Design, Cambridge, UK) and stored using Spike2 software (Cambridge Electronic Design, Cambridge, UK). Using the Spike2 software, data was further processed using a band pass filtered (200-700Hz, 40Hz transition gap) then rectified and smoothed (20ms). Using the Spike2 software, the ECGdelete 02.s2s script is used to remove heart artifact, when present, from the ENG and EMG recordings.

We evaluated swallows that were trigged by injection of water into the oral cavity. *Total swallow duration* was determined by the onset of the submental complex to the termination of the laryngeal complex EMG activity. *Submental complex (SC)* was determined by the onset to the termination of the submental complex EMG activity. *Laryngeal complex (LC)* was determined by the onset to the termination of the laryngeal complex EMG activity. *Swallow sequence* was calculated as the time difference between the peak activation of the laryngeal and submental complex EMG activity. *Schluckatmung* duration was determined by the onset to the termination of the diaphragm EMG activity during a swallow. *Diaphragm inter-burst interval (IBI)* was calculated as the offset of the diaphragm EMG activity to the onset of the subsequent breath. *Inspiratory delay* was calculated as the offset of the swallow-related laryngeal complex EMG activity to the onset of the subsequent breath. Duration and amplitude of each nerve and muscle was determined by the onset to the termination of that respective nerve/muscle activity during swallow. All durations are reported in milliseconds (ms) unless otherwise stated.

Line graphs of swallow frequency in relation to inspiration were created by the phase of breathing in which swallow occurred in, calculated as the onset of inspiration to the onset of swallow divided by the respiratory cycle duration and plotted against the number of swallows that occurred within the 1/10 binned respiratory phase, (*swallow onset: insp onset)*. Swallow was also plotted in relation to the peak activation of the diaphragm as a duration with zero equaling the peak of the inspiratory related diaphragm activity, (*swallow onset: insp peak)*. All data are expressed as mean ± standard deviation (SD), unless otherwise noted. Statistical analyses were performed using GraphPad Prism 10 (GraphPad Software®, Inc. La Jolla, USA). Differences were considered significant at *p* ≤ 0.05. Investigators were not blinded during analysis. Sample sizes were chosen on the basis of previous studies.

## Histology

Upon completion of experiments, animals underwent deep anesthesia using 4% isoflurane in 100% oxygen and were subsequently perfused via the ascending aorta with 20 ml of phosphate buffered saline (PBS; pH 7.4), followed by 20 ml of 4% phosphate-buffered paraformaldehyde (0.1 M; pH 7.4), obtained from Electron Microscopy Sciences, Fort Washington, PA. The brains were then extracted and immersed in the perfusion fixative for 4 hours at 4 °C, followed by an additional 8 hours in 20% sucrose solution. Coronal brain sections (25 μm) were obtained using a cryostat and stored in a cryoprotectant solution at −20°C (consisting of 20% glycerol and 30% ethylene glycol in 50 ml phosphate buffer, pH 7.4) prior to histological processing. All histochemical procedures were conducted on free-floating sections.

IBA-1 was identified using a rabbit anti-IBA-1 antibody (19-19741; Fujifilm Wako Chemicals USA Corp) at a dilution of 1:1000. The antibody was diluted in phosphate buffer (PB) containing 2% normal donkey serum (017-000-121, Jackson ImmunoResearch Laboratories) and 0.3% Triton X-100, and incubated for 24 hours. Following multiple rinses, the sections were incubated with donkey anti-rabbit Alexa 594 (711-585-152, Jackson ImmunoResearch Laboratories) at a dilution of 1:400 in PB for 4 hours at room temperature. After further rinsing in PB, the sections were mounted sequentially in rostrocaudal order onto gelatin-coated slides and cover-slipped with DPX mounting medium (06522; Sigma Aldrich) for histological examination.

For cell counting, imaging, and data analysis, a VS120-S6-W Virtual Slide Scanner (Olympus) was utilized to scan all sections, with images captured using a Nikon DS-Fi3 color camera. To mitigate potential biases, photomicrography and counting were performed by an anatomist blinded to the experimental conditions. Cell counting was conducted using Image J (version 1.41; National Institutes of Health, Bethesda, MD), while line drawings were created using Canvas software (ACD Systems, Victoria, Canada, v. 9.0). Brain sections were analyzed in a one-in-two series of 25-µm sections per mouse, spaced 50 µm apart. The analyzed area was delineated based on previous reports (Anderson et al., 2016). Counts were performed bilaterally, averaged, and reported as mean ± standard deviation (SD). Section alignment was referenced to a standard section, as previously described (Anderson et al., 2016), and based on Paxinos and Franklin (Paxinos & Franklin, 2019). Image J software (W.S. Rasband/National Institutes of Health, USA, http://rsb.info.nih.gov/ij) was also utilized for morphological analysis (fractal and skeleton) of microglia labeled in 40x magnification photomicrographs. The total number of IBA-1 labeled microglia was counted using a cell counter plugin in 4 photomicrographs per analyzed region (VRC, PiCo and NTS). For morphological analysis, binary photomicrographs were first converted into skeletonized images using the Analyze Skeleton Plugin to quantify the number of endpoints and the size of each cell branch (process length).

All collected data were normalized by the total number of cells in each slice. Additionally, 6 cells in binary images were chosen per animal using a rectangle tool (width: 215 µm; height: 175 µm). These isolated cells were converted into outline cells and analyzed using the FracLac plugin. This plugin calculates various parameters including fractal dimension (regression slope (In(N)/ In(Ɛ)), where N is the number of pixels and Ɛ is a particular scale), cell density (number of pixels within cell outline/ area of convex hull), span ratio (convex hull eclipse longest length/ convex hull eclipse longest width), lacunarity (variation in pixel density), and circularity (circularity of the convex hull (4π×area⁄perimeter^2^, for a circle it is 1). Fractal dimension reflects cell complexity, lacunarity indicates cell heterogeneity, density corresponds to cell size, span ratio reflects cell elongation, and circularity indicates cell circularity (Forsberg et al., 2017; Lima-Silveira et al., 2019; Morrison et al., 2017). The data are reported as the mean ± standard deviation. Statistical analysis was performed using GraphPad Prism10. An ordinary one-way ANOVA followed by Tukey’s multiple comparisons test was used.

## Results

### Variable baseline swallow motor pattern is observed in Vglut2, not Gad2-, Ndufs4-KO mice

Swallow related muscle activation pattern must be precisely shaped and timed for safe and efficient bolus transport. Whether Ndufs4 KO mouse models of LS recapitulate swallowing defects seen in LS patients is unknown. To address this knowledge gap, simultaneous submental, laryngeal, and diaphragmatic EMG, along with hypoglossal, and vagal ENG were recorded in mice exposed to room air to evaluate baseline swallow and breathing activity as well as their coordination in control (N=10),Vglut2- (N=13), and Gad2- (N=5) Ndufs4-KO mice within the age range of peak phenotypic severity (11, 28). The control mice were 111 ± 5 days old and weighed 26 ± 2 g. As expected, based on previous characterization of the mutant phenotypes (11), the Vglut2-Ndufs4-KO mice exhibited a significantly lower body weight (19 ± 2 vs 26 ± 2g, p< 0.0001) at comparable age (109 ± 18 vs 111 ± 5, p= 0.76). On the other end, since Gad2-Ndufs4-KO mice die at a younger age without detectable loss in body weight, mice of slightly younger age (71 ± 7 vs 111 ± 5, p= 0.001) but with comparable body weight (23 ± 5 vs 26 ± 2g, p= 0.23) to controls were used (Table 1).

**Table 1.**
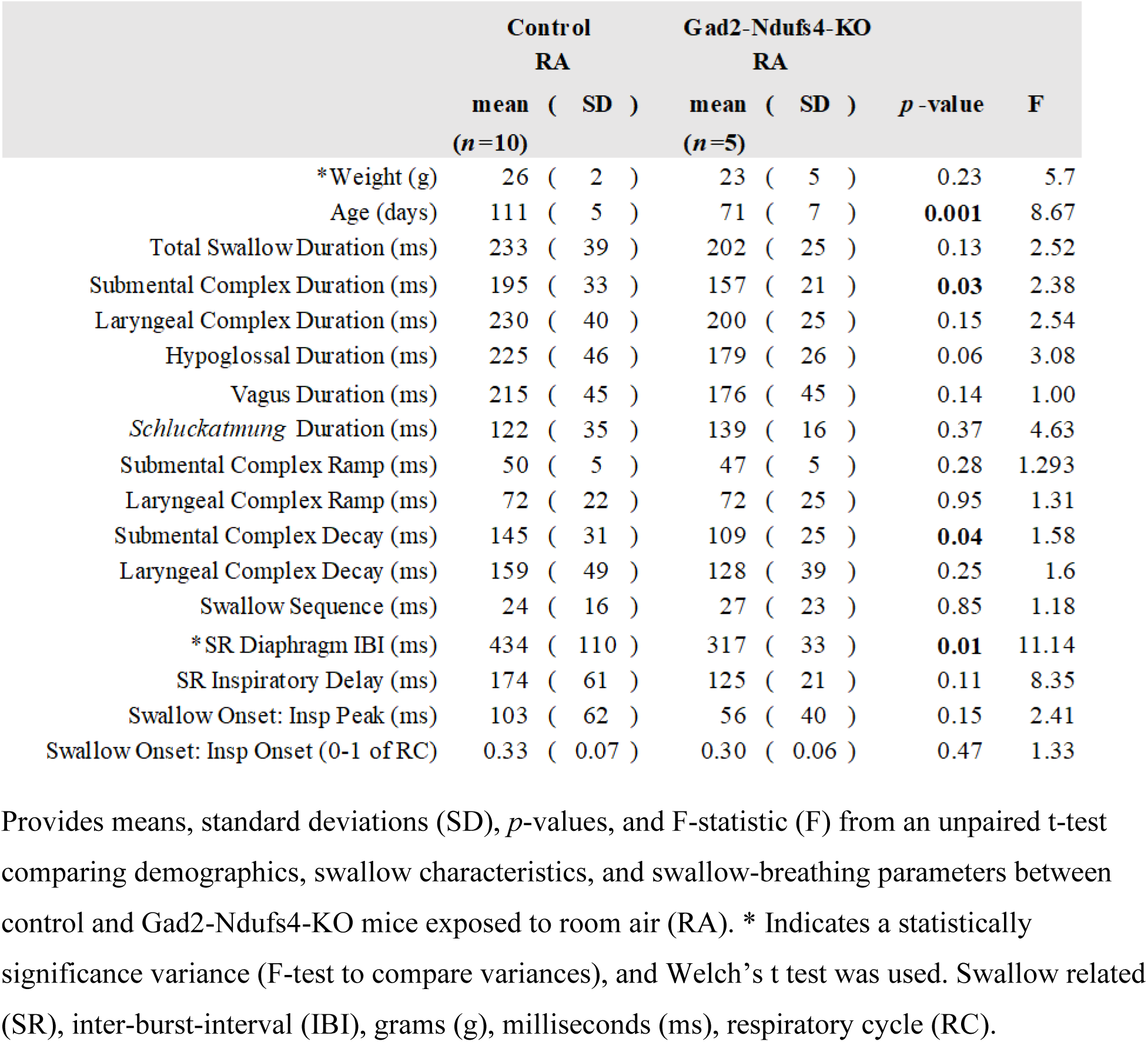
Characteristics of control and Gad2-Ndufs4-KO mice exposed to room air.

EMG traces of water-evoked swallow motor pattern revealed a decrease in submental complex duration and activation decay in the Gad2-Ndufs4-KO mice exposed to room air (Table 1). Whereas, in Vglut2-Ndufs4-KO mice exposed to the same condition, the duration of laryngeal adduction (indicated by laryngeal complex (LC) activity) (311 ± 131ms vs 230 ± 40ms, *p* = 0.05) and ramping of the muscle activity (108 ± 37ms vs 72 ± 22ms, *p* = 0.01) were significantly longer than in control mice (paired t test with Welch’s correction, Table 2). In addition, a putative pharyngeal clearing behavior and a likely aspiration reflex (AspR), characterized as a brief but large burst of diaphragm activity, was observed prior to swallow in 4 of the 13 Vglut2-Ndufs4-KO mice. The diaphragm activity was significantly larger in amplitude compared to the prior eupnea burst (182 ± 12 vs 90 ± 13m % max of eupnea, *p* = 0.0001) (figure 1-figure supplement 1). This swallow related AspR was only seen in the Vglut2-Ndufs4-KO mice and never in the control or Gad2-Ndufs4-KO mice.

**Table 2.** Characteristics of control and Vglut2-Ndufs4-KO mice exposed to room air.

|  | Control |  | Vglut2-Ndufs4-KO |  | *p* -value | F |
| --- | --- | --- | --- | --- | --- | --- |
|  | RA |  | RA |  |  |  |
|  | mean ( SD ) |  | mean ( SD ) |  |  |  |
|  | (*n*=10) |  | (*n*=13) |  |  |  |
| *Weight (g) | 26 ( 2 ) |  | 19 ( 2 ) |  | **<0.0001** | 51.86 |
| Age (days) | 111 ( 5 ) |  | 109 ( 18 ) |  | 0.76 | 1.14 |
| *Total Swallow Duration (ms) | 233 ( 39 ) |  | 311 ( 132 ) |  | 0.06 | 11.23 |
| *Submental Complex Duration (ms) | 195 ( 33 ) |  | 232 ( 78 ) |  | 0.15 | 5.68 |
| *Laryngeal Complex Duration (ms) | 230 ( 40 ) |  | 311 ( 131 ) |  | **0.05** | 11.01 |
| Hypoglossal Duration (ms) | 225 ( 46 ) |  | 242 ( 72 ) |  | 0.51 | 2.42 |
| Vagus Duration (ms) | 215 ( 45 ) |  | 235 ( 75 ) |  | 0.48 | 2.74 |
| *Schluckatmung* Duration (ms) | 122 ( 35 ) |  | 165 ( 55 ) |  | 0.08 | 2.42 |
| *Submental Complex Ramp (ms) | 50 ( 5 ) |  | 74 ( 43 ) |  | 0.07 | 67.92 |
| Laryngeal Complex Ramp (ms) | 72 ( 22 ) |  | 108 ( 37 ) |  | **0.01** | 2.95 |
| Submental Complex Decay (ms) | 145 ( 31 ) |  | 157 ( 51 ) |  | 0.52 | 2.75 |
| *Laryngeal Complex Decay (ms) | 159 ( 49 ) |  | 203 ( 115 ) |  | 0.23 | 5.44 |
| Swallow Sequence (ms) | 24 ( 23 ) |  | 33 ( 40 ) |  | 0.55 | 3.02 |
| *SR Diaphragm IBI (ms) | 434 ( 110 ) |  | 926 ( 352 ) |  | **0.0003** | 10.16 |
| *SR Inspiratory Delay (ms) | 174 ( 61 ) |  | 529 ( 315 ) |  | **0.002** | 26.71 |
| *Insp Peak to Swallow Onset (ms) | 103 ( 62 ) |  | 273 ( 244 ) |  | **0.03** | 15.51 |
| Swallow Onset: Insp Onset (0-1 of RC) | 0.33 ( 0.07 ) |  | 0.24 ( 0.08 ) |  | **0.01** | 1.31 |

Provides means, standard deviations (SD), *p*-values, and F-statistic (F) from an unpaired t-test comparing demographics, swallow characteristics, and swallow-breathing parameters between control and Vglut2-Ndufs4-KO mice exposed to room air (RA). \* Indicates a statistically significance variance (F-test to compare variances), and Welch's t test was used. Swallow related (SR), inter-burst-interval (IBI), grams (g), milliseconds (ms), respiratory cycle (RC).

The F test to compare variability in muscle activity that controls swallowing indicated a statistically significant variance between Vglut2-Ndufs4-KO and control mice in the total swallow duration (*F* = 11.23, *p* = 0.001), submental complex (SC) duration (*F* = 5.68, *p* = 0.01), LC duration (*F* = 11.07, *p* = 0.001), SC ramp (*F* = 67.92, *p* < 0.0001), and LC complex decay (*F* = 5.44, *p* = 0.02) by unpaired t test with Welch’s correction. This comparison is visually depicted by overlaying representative swallow traces in figure 1F. Together, these results suggest that major swallowing defects were present exclusively in Vglut2-, not Gad2-, Ndufs4KO mice. These physiological defects arise from an increased variability in swallow motor pattern which likely result in abnormal swallow behavior. This could hinder the ability to properly swallow and ingest food, and subsequently contribute to the decreased body weight and premature death in Vglut2-, but not in the Gad2-Ndufs4-KO mice.

### Swallow production causes disruption to respiratory rhythm regeneration

The coordination of swallow and breathing as well as the regeneration of the respiratory rhythm following a swallow are essential to maintain homeostasis. Our recordings revealed a decreased swallow-related (SR) Diaphragm Inter-Burst Interval (IBI) in Gad-Ndufs4-KO mice (Table 1, Figure 2A). However, the SR -IBI (926 ± 352ms vs 434 ± 110ms, *p* = 0.0003) and inspiratory delay (529 ± 315ms vs 174 ± 61ms, *p* = 0.002) (Figure 2A and B) were significantly longer in duration in Vglut2-Ndufs4-KO mice as compared to control mice. There was also significant variability in SR diaphragm IBI (*F* = 10.16, *p* = 0.002) and inspiratory delay (*F* = 26.71, *p* < 0.0001) indicated by an unpaired t-test with Welch’s correction (Table 2).

To evaluate the recovery of the respiratory rhythm after a swallow, we plotted the average inspiratory delay (breath 1) plus the following 50 respiratory IBI (total 51 breaths) in control, Vglut2-, and Gad2-Ndufs4-KO mice exposed to RA. A two-way mixed effects ANOVA revealed a significant interaction between genotype and IBI (*p* <0.0001, df = 50, F = 5.92). Tukey’s multiple comparison test revealed statistical differences in various breaths from 1-51 between Control and Gad2-Ndufs4-KO mice, which suggested Gad2-Ndufs4-KO mice have a decreased IBI, presumably faster breathing following a swallow. While the post swallow IBI in Vglut2-Ndufs4-KO was significantly longer in duration compared to control from breaths 1-28 (p< 0.03) and did not reach control durations until breath 29 (Figure 2A). There was an increase in the variability of each IBI duration following a swallow in Vglut2-Ndufs4-KO compared to control and Gad2-Ndufs4-KO mice. This is visually depicted in figure 2, which shows representative traces of all 3 genotypes. These analyses reveal abnormalities in swallow behavior and coordination with breathing only in the Vglut2-Ndufs4-KO mice and not the Gad2-Ndufs4-KO mice. This guided us to only focus on the Vglut2-Ndufs4-KO mice.

### Water stimulation induces various types of apneas in Vglut2-Ndufs4-KO mice exposed to room air

In 7 out of the 13 Vglut2-Ndufs4-KO room air mice, three types of water-induced apnea occurred (Figure 3). Apnea was defined as quiescence of the diaphragm two times the average duration of the respiratory IBI across 10 respiratory cycles. Unlike the other parameters examined above, apneas were measured in seconds (s) instead of milliseconds (ms). Type 1, defined as water-induced diaphragm quiescence one time the average IBI duration prior to a swallow (Figure 3B), occurred 3 times in 2 of the 7 mice. Type 2, defined as water-induced apneas with the absence of swallow or any upper airway behavior (Figure 3E), occurred 4 times in 3 of 7 mice. Lastly, Type 3, defined as diaphragm quiescence following water injection with tonic laryngeal activation (Figure 3F), occurred 14 times in 3 out 7 mice. A one-way ANOVA indicated no statistical difference in the duration of water induced apneas across all three types, which ranged from 0.65s to 28s with an average of 4 ± 7s (Figure 3H).

**Figure 3.**
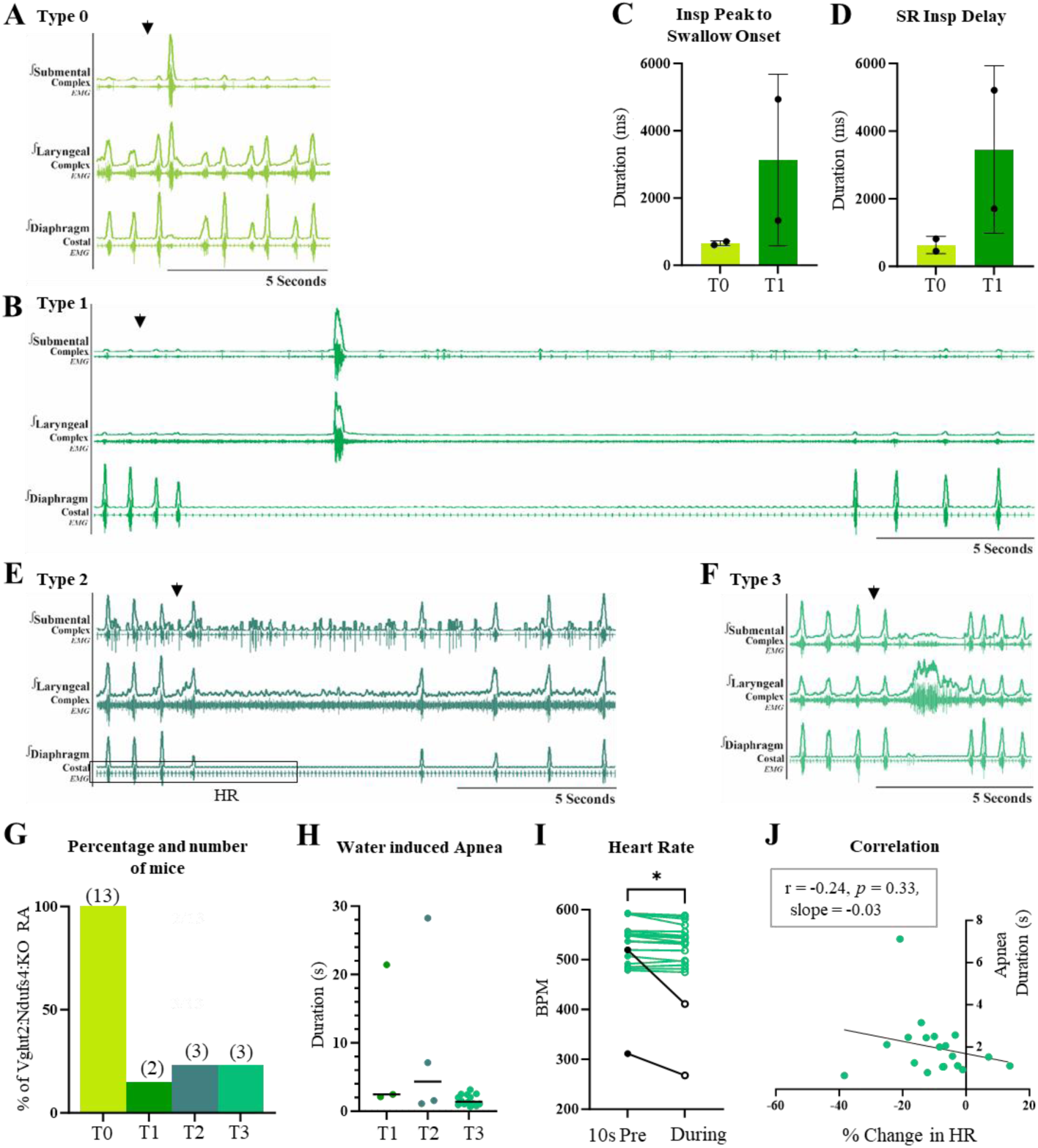
Characteristics of water induced apneas seen only in Vglut2-Ndufs4-KO mice exposed to room air. Apnea was defined as quiescence of the diaphragm two times the average duration of the respiratory IBI across 10 respiratory cycles. A) representative trace of type 0 (T0) where injection of water (arrow) did not induce apnea but a swallow. B) Representative trace of type 1 (T1), defined as a water induced diaphragm quiescence one time the average IBI duration prior to a swallow, occurred 3 times in 2 of the 7 mice. C) Duration of diaphragm quiescence before and D) after a swallow. E) Representative trace of type 2 (T2), defined as water induced apneas with the absence of swallow or any upper airway behavior, occurred 4 times in 3 of 7 mice. F) Representative trace of type 3 (T3), defined as diaphragm quiescence following water injection with tonic laryngeal activation, occurred 14 times in 3 out 7 mice. G) Bar graph representing the percentage of Vglut2-Ndufs4-KO mice exposed to room air that had each type of water induced apnea. H) A one-way ANOVA indicated no statistical difference in the duration of water induced apneas across all three types, which ranged from 0.65s to 28s with an average of 4 ± 7s. I) Average heart rate (HR) 10s prior to and during the water induced apnea. In 19 of the 21 apnea bouts (green), we found a slight but significant decrease in HR during the apnea bout (540 ± 38 bpm, 534 ± 35bpm, *p* = 0.001) compared to 10s prior to apnea with an average percent change of −10 ± 12%. Two of the 21 apneas (black) lasted longer than 20s and had on average a 174% decrease in HR. J) A Pearson correlation found no significant correction between length of apnea and % change in HR.

Measurements of heart rate (HR) in the 10s prior to and during the duration of the apnea were obtained to assess the impact of swallow related apnea on cardiac function. In 19 of the 21 apneas, we found a slight but significant decrease in HR during apnea (540 ± 38 bpm, 534 ± 35bpm, *p* = 0.001) compared to 10s prior to apnea (Figure 3I) with an average percent change of −10 ± 12%. A Pearson correlation found no significant correction between length of apnea and % change in HR (r = −0.24, *p* = 0.33, Figure 3J). Two of the 21 apneas lasted longer than 20s and had on average a 174% decrease in HR. These water-induced apneas never occur in control or Gad2-Ndufs4-KO mice.

### Chronic Hypoxia prevents premature death, swallow motor pattern variability, and maintains respiratory rhythm recovery

Control and Vglut2-Ndufs4-KO mice were exposed to chronic hypoxia (CH) (11% O_2_), prior to symptoms onset at an average age of p45 and 52, respectively, for 6 months. Vglut2-Ndufs4-KO CH mice lived significantly longer than the mice exposed to room air. A Mantel-Cox Log-rank tests indicated the median survival for RA mice is 100 days and 223 days for CH in the Vglut2-Ndufs4-KO mice (*p* <0.0001). It is important to note, experimental protocol for physiological assessments required alive, yet anesthetized mice, which resulted in artificially picking their end date, see methods for criteria. A two-way mixed-effects ANOVA revealed a significant interaction between genotype and measured swallow characteristics (*p* <0.0001, df = 1.3, F = 54.20) (Table 3). Tukey’s multiple comparison test revealed Vglut2-Ndufs4-KO CH mice weighed significantly more than Vglut2-Ndufs4-KO RA mice (24 ± 3g, 19 ± 2g, *p* = 0.003), but there was no difference compared to the control mice exposed to CH. Vglut2-Ndufs4-KO CH mice had a significant decrease in total swallow (187 ± 48ms, 311 ± 132ms, *p* = 0.03), submental complex (147 ± 43ms, 232 ± 78ms, *p* = 0.01), and laryngeal complex duration (183 ± 43ms, 311 ± 131ms, *p* = 0.02) compared to RA, but there is no change compared to control CH mice (Figure 4D, Table 3). Unlike the Vglut2-Ndufs4-KO RA mice (Figure 1F), there is no significant change in duration or variance between control CH and Vglut2-Ndufs4-KO CH mice in all measured parameters, visually depicted by overlaid representative traces of swallow in figure 4C (Table 3). AspR and apneas seen in the RA mice were never observed in the CH mice. We suggest the decreased variability in CH exposed mice results in normal swallow activity and proper ingestion, without risking airway protection.

**Figure 4.**
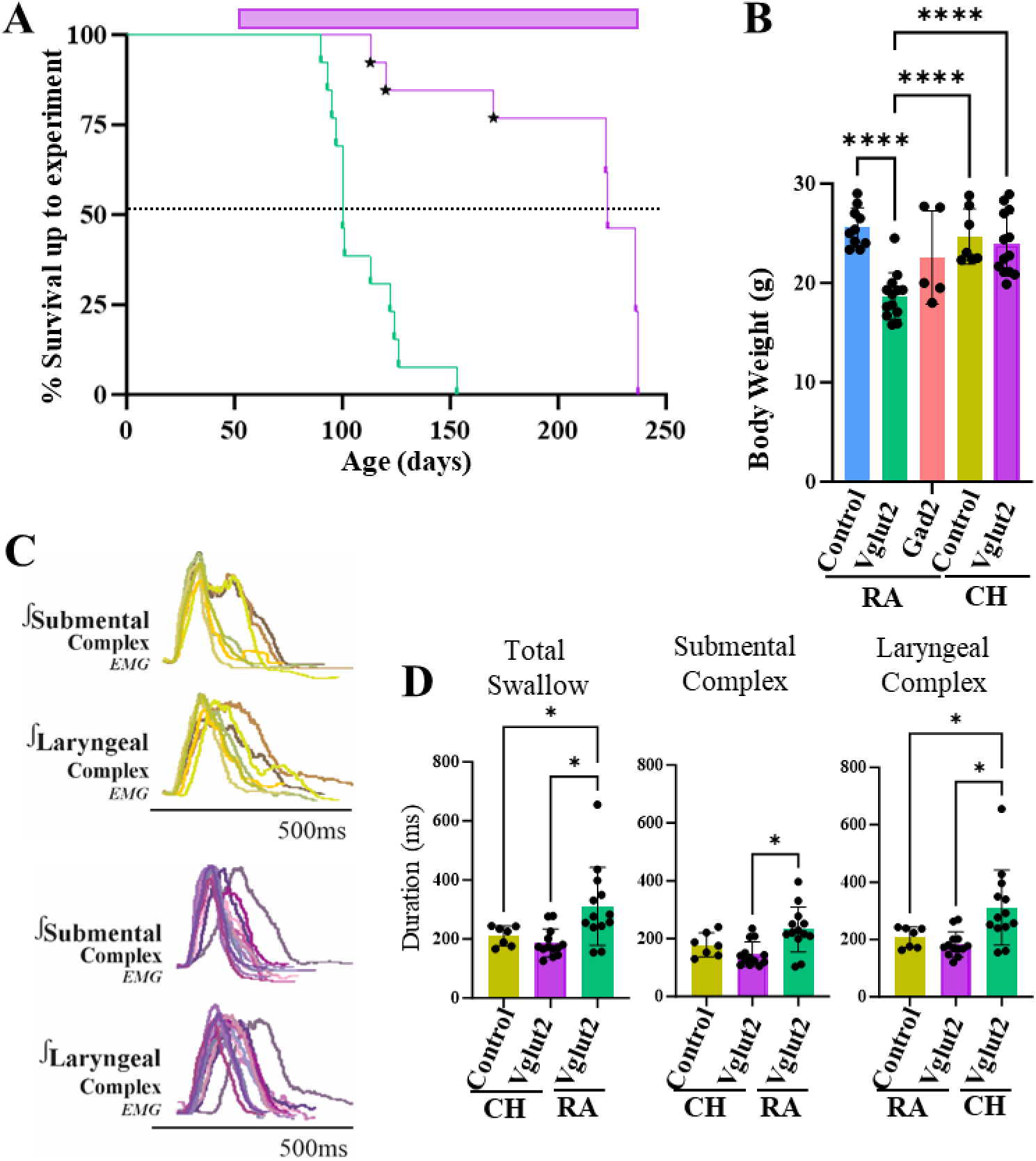
Exposure to chronic hypoxia (CH) increases life expectancy, maintains body weight, and prevents alteration and variability in swallow motor pattern. A) A Mantel-Cox Log-rank test revealed a significant increase in survival for Vglut2-Ndufs4-KO mice exposed to CH (*p* <0.0001). Purple bar at the top indicated time in CH. B) We found no significant difference in body weight of control mice exposed to room air (RA), control mice exposed to CH, or Vglut2-Ndufs4-KO mice exposed to CH. C) CH prevents variability, seen in RA (Figure 1F) of swallow motor patterning with no change in ramp or decay of the submental and laryngeal complex activity. D) CH prevents the increase in total swallow, submental and laryngeal complex duration seen in Vglut2-Ndufs4-KO mice exposed to room air.

**Table 3.** Characteristics of control and Vglut2-Ndufs4-KO mice exposed to room air and chronic hypoxia.

|  | Control<br>CH | Vglut2-Ndufs4-KO<br>CH | Vglut2-Ndufs4-KO<br>RA | *p*-value |
| --- | --- | --- | --- | --- |
|  | mean ( SD )<br>(*n*=7) | mean ( SD )<br>(*n*=13) | mean ( SD )<br>(*n*=13) |  |
| Weight (g) | 25 ( 3 ) | 24 ( 3 ) | 19 ( 2 ) |  |
|  |  |  | *Con* vs *Vglut2 RA* | **0.0003** |
|  |  |  | *Con* vs *Vglut2 CH* | 0.87 |
|  |  |  | *Vglut2 RA* vs *Vglut2 CH* | **0.003** |
| Age (days) | 165 ( 3 ) | 209 ( 45 ) | 109 ( 18 ) |  |
|  |  |  | *Con CH* vs *Vglut2 RA* | **<0.0001** |
|  |  |  | *Con CH* vs *Vglut2 CH* | **0.002** |
|  |  |  | *Vglut2 RA* vs *Vglut2 CH* | **<0.0001** |
| Total Swallow Duration (ms) | 211 ( 33 ) | 187 ( 48 ) | 311 ( 132 ) |  |
|  |  |  | *Con CH* vs *Vglut2 RA* | **0.05** |
|  |  |  | *Con CH* vs *Vglut2 CH* | 0.62 |
|  |  |  | *Vglut2 RA* vs *Vglut2 CH* | **0.03** |
| Submental Complex Duration (ms) | 178 ( 41 ) | 147 ( 43 ) | 232 ( 78 ) |  |
|  |  |  | *Con CH* vs *Vglut2 RA* | 0.42 |
|  |  |  | *Con CH* vs *Vglut2 CH* | 0.46 |
|  |  |  | *Vglut2 RA* vs *Vglut2 CH* | **0.01** |
| Laryngeal Complex Duration (ms) | 208 ( 36 ) | 183 ( 43 ) | 311 ( 131 ) |  |
|  |  |  | *Con CH* vs *Vglut2 RA* | **0.04** |
|  |  |  | *Con CH* vs *Vglut2 CH* | 0.60 |
|  |  |  | *Vglut2 RA* vs *Vglut2 CH* | **0.02** |
| Hypoglossal Duration (ms) | 194 ( 42 ) | 164 ( 34 ) | 242 ( 72 ) |  |
|  |  |  | *Con CH* vs *Vglut2 RA* | 0.43 |
|  |  |  | *Con CH* vs *Vglut2 CH* | 0.48 |
|  |  |  | *Vglut2 RA* vs *Vglut2 CH* | **0.006** |
| Vagus Duration (ms) | 186 ( 43 ) | 161 ( 33 ) | 235 ( 75 ) |  |
|  |  |  | *Con CH* vs *Vglut2 RA* | 0.49 |
|  |  |  | *Con CH* vs *Vglut2 CH* | 0.52 |
|  |  |  | *Vglut2 RA* vs *Vglut2 CH* | **0.01** |
| Schluckatmung Duration (ms) | 121 ( 11 ) | 93 ( 27 ) | 165 ( 55 ) |  |
|  |  |  | *Con CH* vs *Vglut2 RA* | 0.37 |
|  |  |  | *Con CH* vs *Vglut2 CH* | 0.22 |
|  |  |  | *Vglut2 RA* vs *Vglut2 CH* | 0.29 |
| Submental Complex Ramp (ms) | 45 ( 4 ) | 47 ( 9 ) | 74 ( 43 ) |  |
|  |  |  | *Con CH* vs *Vglut2 RA* | 0.30 |
|  |  |  | *Con CH* vs *Vglut2 CH* | 0.92 |
|  |  |  | *Vglut2 RA* vs *Vglut2 CH* | 0.10 |
| Laryngeal Complex Ramp (ms) | 65 ( 24 ) | 67 ( 14 ) | 108 ( 37 ) |  |
|  |  |  | *Con CH* vs *Vglut2 RA* | 0.11 |
|  |  |  | *Con CH* vs *Vglut2 CH* | 0.98 |
|  |  |  | *Vglut2 RA* vs *Vglut2 CH* | **0.004** |
| Submental Complex Decay (ms) | 133 ( 42 ) | 100 ( 36 ) | 157 ( 51 ) |  |
|  |  |  | *Con CH* vs *Vglut2 RA* | 0.63 |
|  |  |  | *Con CH* vs *Vglut2 CH* | 0.41 |
|  |  |  | *Vglut2 RA* vs *Vglut2 CH* | **0.01** |
| Laryngeal Complex Decay (ms) | 143 ( 35 ) | 116 ( 42 ) | 203 ( 115 ) |  |
|  |  |  | *Con CH* vs *Vglut2 RA* | 0.10 |
|  |  |  | *Con CH* vs *Vglut2 CH* | 0.42 |
|  |  |  | *Vglut2 RA* vs *Vglut2 CH* | 0.09 |
| Swallow Sequence (ms) | 23 ( 25 ) | 24 ( 11 ) | 33 ( 40 ) |  |
|  |  |  | *Con CH* vs *Vglut2 RA* | 0.82 |
|  |  |  | *Con CH* vs *Vglut2 CH* | 1.00 |
|  |  |  | *Vglut2 RA* vs *Vglut2 CH* | 0.67 |
| SR Diaphragm IBI (ms) | 431 ( 125 ) | 460 ( 160 ) | 926 ( 352 ) |  |
|  |  |  | *Con CH* vs *Vglut2 RA* | **0.01** |
|  |  |  | *Con CH* vs *Vglut2 CH* | 0.92 |
|  |  |  | *Vglut2 RA* vs *Vglut2 CH* | **0.01** |
| SR Inspiratory Delay (ms) | 189 ( 84 ) | 268 ( 139 ) | 529 ( 315 ) |  |
|  |  |  | *Con CH* vs *Vglut2 RA* | **0.02** |
|  |  |  | *Con CH* vs *Vglut2 CH* | 0.22 |
|  |  |  | *Vglut2 RA* vs *Vglut2 CH* | 0.11 |
| Swallow Onset: Insp Peak (ms) | 106 ( 63 ) | 71 ( 34 ) | 273 ( 244 ) |  |
|  |  |  | *Con CH* vs *Vglut2 RA* | 0.26 |
|  |  |  | *Con CH* vs *Vglut2 CH* | 0.30 |
|  |  |  | *Vglut2 RA* vs *Vglut2 CH* | **0.03** |
| Swallow Onset: Insp Onset (0-1 of RC) | 0.32 ( 0.04 ) | 0.25 ( 0.05 ) | 0.24 ( 0.08 ) |  |
|  |  |  | *Con CH* vs *Vglut2 RA* | 0.08 |
|  |  |  | *Con CH* vs *Vglut2 CH* | **0.01** |
|  |  |  | *Vglut2 RA* vs *Vglut2 CH* | 0.89 |

Provides means, standard deviations (SD), and *p*-values from a two-way mixed effects ANOVA using Tukey's multiple comparison test comparing demographics, swallow characteristics and swallow-breathing parameters between control (con) and Vglut2-Ndufs4-KO mice exposed to chronic hypoxia (CH), and Vglut2-Ndufs4-KO mice exposed to room air (RA). Swallow related (SR), inter-burst-interval (IBI), grams (g), milliseconds (ms), respiratory cycle (RC).

**Table 4.** Sex-specific characteristics of control and Gad2-Ndufs4-KO mice exposed to room air.

| Gad2-Ndufs4-KO | Male (n=3) |  | Female (n=2) |  | p-value | F | t | df |
| --- | --- | --- | --- | --- | --- | --- | --- | --- |
| RA | mean | ( SD ) | mean | ( SD ) |  |  |  |  |
| Weight (g) | 25 | ( 4 ) | 19 | ( 1 ) | 0.15 | - | 1.90 | 3 |
| Age (days) | 73 | ( 7 ) | 69 | ( 7 ) | 0.61 | - | 0.56 | 3 |
| Total Swallow Duration (ms) | 208 | ( 19 ) | 192 | ( 38 ) | 0.54 | - | 0.69 | 3 |
| Submental Complex Duration (ms) | 154 | ( 27 ) | 160 | ( 17 ) | 0.81 | - | 0.26 | 3 |
| Laryngeal Complex Duration (ms) | 207 | ( 20 ) | 191 | ( 37 ) | 0.57 | - | 0.64 | 3 |
| Hypoglossal Duration (ms) | 189 | ( 24 ) | 164 | ( 31 ) | 0.38 | - | 1.03 | 3 |
| Vagus Duration (ms) | 181 | ( 58 ) | 167 | ( 35 ) | 0.79 | - | 0.29 | 3 |
| *Schluckatmung* Duration (ms) | 143 | ( 18 ) | 129 | ( - ) | 0.56 | - | 0.69 | 2 |
| Submental Complex Ramp (ms) | 50 | ( 4 ) | 43 | ( 2 ) | 0.12 | - | 2.19 | 3 |
| Laryngeal Complex Ramp (ms) | 82 | ( 30 ) | 59 | ( 3 ) | 0.39 | - | 1.01 | 3 |
| Submental Complex Decay (ms) | 105 | ( 30 ) | 117 | ( 19 ) | 0.66 | - | 0.49 | 3 |
| Laryngeal Complex Decay (ms) | 125 | ( 50 ) | 132 | ( 34 ) | 0.88 | - | 0.17 | 3 |
| Swallow Sequence (ms) | 33 | ( 33 ) | 16 | ( 7 ) | 0.54 | - | 0.69 | 3 |
| SR Diaphragm IBI (ms) | 310 | ( 38 ) | 328 | ( 33 ) | 0.62 | - | 0.55 | 3 |
| SR Inspiratory Delay (ms) | 114 | ( 19 ) | 141 | ( 14 ) | 0.19 | - | 1.67 | 3 |
| Swallow Onset: Insp Peak (ms) | 53 | ( 52 ) | 61 | ( 28 ) | 0.84 | - | 0.21 | 3 |
| Swallow Onset: Insp Onset (0-1 of RC) | 0.28 | ( 0.07 ) | 0.34 | ( 0.01 ) | 0.36 | - | 1.08 | 3 |

Provides means, standard deviations (SD), *p*-values, F-statistic (F), t-statistics (t), and degrees of freedom (df) from an unpaired t-test comparing demographics, swallow characteristics and swallow-breathing parameters between male and female Gad2-Ndufs4-KO mice exposed to room air (RA). Swallow related (SR), inter-burst-interval (IBI), grams (g), milliseconds (ms), respiratory cycle (RC).

**Table 5.**
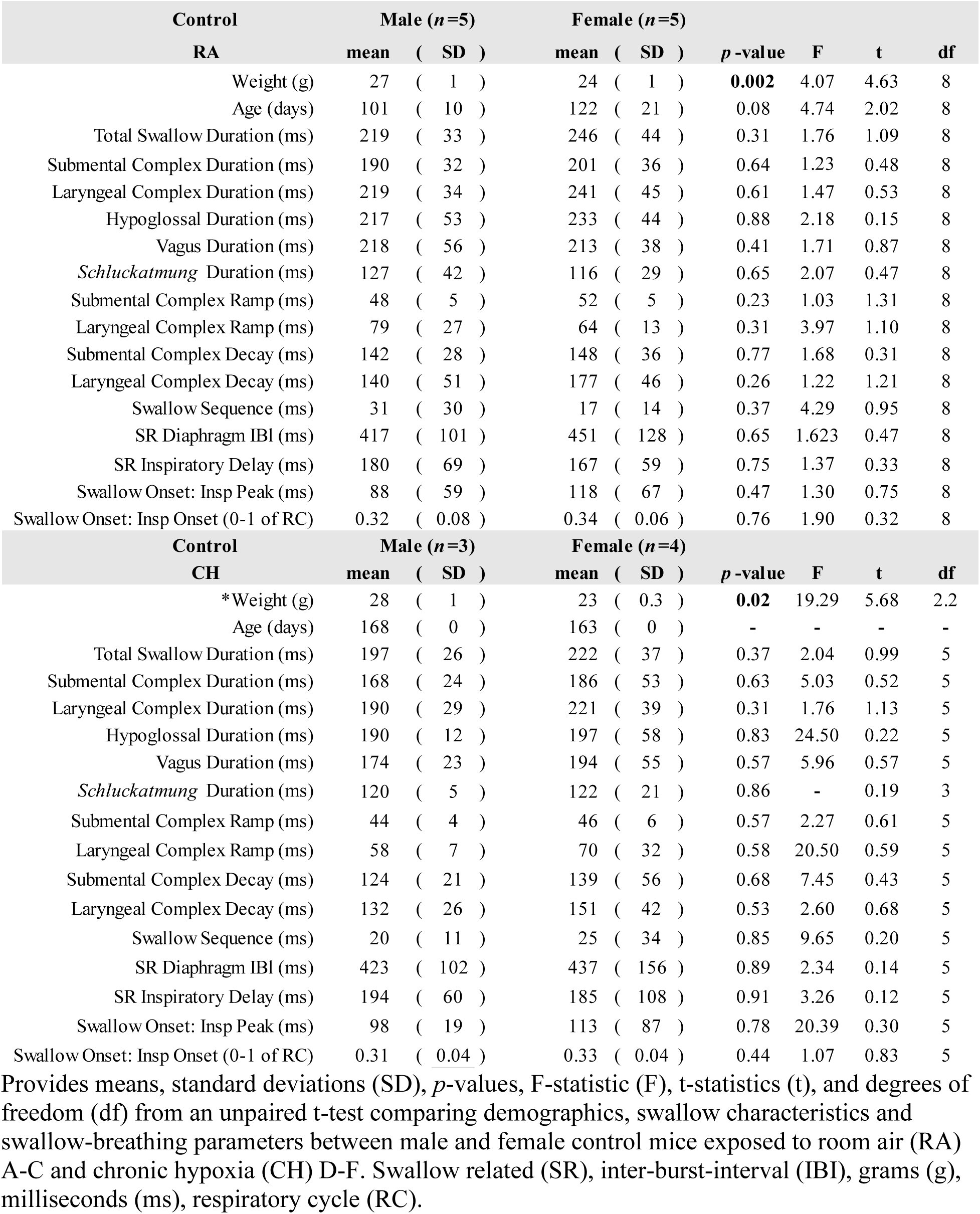
Sex-specific characteristics of control mice exposed to room air and chronic hypoxia.

| Control | Male (n=5) |  | Female (n=5) |  | p -value | F | t | df |
| --- | --- | --- | --- | --- | --- | --- | --- | --- |
| RA | mean | ( SD ) | mean | ( SD ) |  |  |  |  |
| Weight (g) | 27 | ( 1 ) | 24 | ( 1 ) | **0.002** | 4.07 | 4.63 | 8 |
| Age (days) | 101 | ( 10 ) | 122 | ( 21 ) | 0.08 | 4.74 | 2.02 | 8 |
| Total Swallow Duration (ms) | 219 | ( 33 ) | 246 | ( 44 ) | 0.31 | 1.76 | 1.09 | 8 |
| Submental Complex Duration (ms) | 190 | ( 32 ) | 201 | ( 36 ) | 0.64 | 1.23 | 0.48 | 8 |
| Laryngeal Complex Duration (ms) | 219 | ( 34 ) | 241 | ( 45 ) | 0.61 | 1.47 | 0.53 | 8 |
| Hypoglossal Duration (ms) | 217 | ( 53 ) | 233 | ( 44 ) | 0.88 | 2.18 | 0.15 | 8 |
| Vagus Duration (ms) | 218 | ( 56 ) | 213 | ( 38 ) | 0.41 | 1.71 | 0.87 | 8 |
| *Schluckatmung* Duration (ms) | 127 | ( 42 ) | 116 | ( 29 ) | 0.65 | 2.07 | 0.47 | 8 |
| Submental Complex Ramp (ms) | 48 | ( 5 ) | 52 | ( 5 ) | 0.23 | 1.03 | 1.31 | 8 |
| Laryngeal Complex Ramp (ms) | 79 | ( 27 ) | 64 | ( 13 ) | 0.31 | 3.97 | 1.10 | 8 |
| Submental Complex Decay (ms) | 142 | ( 28 ) | 148 | ( 36 ) | 0.77 | 1.68 | 0.31 | 8 |
| Laryngeal Complex Decay (ms) | 140 | ( 51 ) | 177 | ( 46 ) | 0.26 | 1.22 | 1.21 | 8 |
| Swallow Sequence (ms) | 31 | ( 30 ) | 17 | ( 14 ) | 0.37 | 4.29 | 0.95 | 8 |
| SR Diaphragm IBI (ms) | 417 | ( 101 ) | 451 | ( 128 ) | 0.65 | 1.623 | 0.47 | 8 |
| SR Inspiratory Delay (ms) | 180 | ( 69 ) | 167 | ( 59 ) | 0.75 | 1.37 | 0.33 | 8 |
| Swallow Onset: Insp Peak (ms) | 88 | ( 59 ) | 118 | ( 67 ) | 0.47 | 1.30 | 0.75 | 8 |
| Swallow Onset: Insp Onset (0-1 of RC) | 0.32 | ( 0.08 ) | 0.34 | ( 0.06 ) | 0.76 | 1.90 | 0.32 | 8 |
| Control | Male (n=3) |  | Female (n=4) |  | p -value | F | t | df |
| CH | mean | ( SD ) | mean | ( SD ) |  |  |  |  |
| *Weight (g) | 28 | ( 1 ) | 23 | ( 0.3 ) | **0.02** | 19.29 | 5.68 | 2.2 |
| Age (days) | 168 | ( 0 ) | 163 | ( 0 ) | - | - | - | - |
| Total Swallow Duration (ms) | 197 | ( 26 ) | 222 | ( 37 ) | 0.37 | 2.04 | 0.99 | 5 |
| Submental Complex Duration (ms) | 168 | ( 24 ) | 186 | ( 53 ) | 0.63 | 5.03 | 0.52 | 5 |
| Laryngeal Complex Duration (ms) | 190 | ( 29 ) | 221 | ( 39 ) | 0.31 | 1.76 | 1.13 | 5 |
| Hypoglossal Duration (ms) | 190 | ( 12 ) | 197 | ( 58 ) | 0.83 | 24.50 | 0.22 | 5 |
| Vagus Duration (ms) | 174 | ( 23 ) | 194 | ( 55 ) | 0.57 | 5.96 | 0.57 | 5 |
| *Schluckatmung* Duration (ms) | 120 | ( 5 ) | 122 | ( 21 ) | 0.86 | - | 0.19 | 3 |
| Submental Complex Ramp (ms) | 44 | ( 4 ) | 46 | ( 6 ) | 0.57 | 2.27 | 0.61 | 5 |
| Laryngeal Complex Ramp (ms) | 58 | ( 7 ) | 70 | ( 32 ) | 0.58 | 20.50 | 0.59 | 5 |
| Submental Complex Decay (ms) | 124 | ( 21 ) | 139 | ( 56 ) | 0.68 | 7.45 | 0.43 | 5 |
| Laryngeal Complex Decay (ms) | 132 | ( 26 ) | 151 | ( 42 ) | 0.53 | 2.60 | 0.68 | 5 |
| Swallow Sequence (ms) | 20 | ( 11 ) | 25 | ( 34 ) | 0.85 | 9.65 | 0.20 | 5 |
| SR Diaphragm IBI (ms) | 423 | ( 102 ) | 437 | ( 156 ) | 0.89 | 2.34 | 0.14 | 5 |
| SR Inspiratory Delay (ms) | 194 | ( 60 ) | 185 | ( 108 ) | 0.91 | 3.26 | 0.12 | 5 |
| Swallow Onset: Insp Peak (ms) | 98 | ( 19 ) | 113 | ( 87 ) | 0.78 | 20.39 | 0.30 | 5 |
| Swallow Onset: Insp Onset (0-1 of RC) | 0.31 | ( 0.04 ) | 0.33 | ( 0.04 ) | 0.44 | 1.07 | 0.83 | 5 |

Provides means, standard deviations (SD), *p*-values, F-statistic (F), t-statistics (t), and degrees of freedom (df) from an unpaired t-test comparing demographics, swallow characteristics and swallow-breathing parameters between male and female control mice exposed to room air (RA) A-C and chronic hypoxia (CH) D-F. Swallow related (SR), inter-burst-interval (IBI), grams (g), milliseconds (ms), respiratory cycle (RC).

**Table 6.** Sex-specific characteristics of Vglut2-Ndufs4-KO mice exposed to room air and chronic hypoxia.

| Vglut2-Ndufs4-KO |  | Male (n=7) |  | Female (n=6) |  | p-value | F | t | df |
| --- | --- | --- | --- | --- | --- | --- | --- | --- | --- |
| RA |  | mean | ( SD ) | mean | ( SD ) |  |  |  |  |
|  | Weight (g) | 20 | ( 3 ) | 17 | ( 1 ) | 0.08 | 2.90 | 1.94 | 11 |
|  | Age (days) | 112 | ( 21 ) | 106 | ( 15 ) | 0.57 | 2.13 | 0.59 | 11 |
|  | Total Swallow Duration (ms) | 306 | ( 96 ) | 316 | ( 175 ) | 0.90 | 3.33 | 0.13 | 11 |
|  | Submental Complex Duration (ms) | 236 | ( 85 ) | 227 | ( 77 ) | 0.85 | 1.23 | 0.20 | 11 |
|  | Laryngeal Complex Duration (ms) | 306 | ( 94 ) | 317 | ( 175 ) | 0.88 | 3.48 | 0.15 | 11 |
|  | Hypoglossal Duration (ms) | 245 | ( 82 ) | 239 | ( 67 ) | 0.90 | 1.51 | 0.13 | 11 |
|  | Vagus Duration (ms) | 240 | ( 86 ) | 228 | ( 67 ) | 0.78 | 1.65 | 0.28 | 11 |
|  | *Schluckatmung* Duration (ms) | 157 | ( 36 ) | 175 | ( 82 ) | 0.69 | 5.15 | 0.42 | 5 |
|  | Submental Complex Ramp (ms) | 78 | ( 44 ) | 70 | ( 45 ) | 0.74 | 1.03 | 0.33 | 11 |
|  | Laryngeal Complex Ramp (ms) | 103 | ( 32 ) | 114 | ( 44 ) | 0.61 | 1.91 | 0.52 | 11 |
|  | Submental Complex Decay (ms) | 158 | ( 51 ) | 157 | ( 56 ) | 0.98 | 1.18 | 0.24 | 11 |
|  | Laryngeal Complex Decay (ms) | 203 | ( 85 ) | 204 | ( 152 ) | 0.99 | 3.17 | 0.01 | 11 |
|  | Swallow Sequence (ms) | 25 | ( 44 ) | 42 | ( 38 ) | 0.45 | 1.34 | 0.78 | 11 |
|  | SR Diaphragm IBI (ms) | 1143 | ( 695 ) | 980 | ( 443 ) | 0.63 | 2.46 | 0.49 | 11 |
|  | SR Inspiratory Delay (ms) | 626 | ( 463 ) | 544 | ( 409 ) | 0.74 | 1.28 | 0.74 | 11 |
|  | Swallow Onset: Insp Peak (ms) | 355 | ( 296 ) | 177 | ( 130 ) | 0.20 | 5.20 | 1.36 | 11 |
|  | Swallow Onset: Insp Onset (0-1 of RC) | 0.23 | ( 0.07 ) | 0.25 | ( 0.09 ) | 0.66 | 1.45 | 0.46 | 11 |
| Vglut2-Ndufs4-KO |  | Male (n=5) |  | Female (n=8) |  | p-value | F | t | df |
| CH |  | mean | ( SD ) | mean | ( SD ) |  |  |  |  |
|  | Weight (g) | 27 | ( 2 ) | 22 | ( 1 ) | **<0.0001** | 1.50 | 5.98 | 11 |
|  | *Age (days) | 228 | ( 7 ) | 197 | ( 54 ) | 0.15 | 52.64 | 1.60 | 7.4 |
|  | Total Swallow Duration (ms) | 159 | ( 24 ) | 204 | ( 51 ) | 0.09 | 4.43 | 1.83 | 11 |
|  | Submental Complex Duration (ms) | 128 | ( 20 ) | 158 | ( 50 ) | 0.22 | 6.09 | 1.29 | 11 |
|  | Laryngeal Complex Duration (ms) | 156 | ( 25 ) | 200 | ( 45 ) | 0.08 | 3.12 | 1.96 | 11 |
|  | Hypoglossal Duration (ms) | 156 | ( 23 ) | 169 | ( 40 ) | 0.51 | 3.01 | 0.68 | 11 |
|  | Vagus Duration (ms) | 152 | ( 21 ) | 166 | ( 39 ) | 0.48 | 3.66 | 0.73 | 11 |
|  | *Schluckatmung* Duration (ms) | 72 | ( 14 ) | 109 | ( 23 ) | 0.06 | 2.81 | 2.43 | 5 |
|  | *Submental Complex Ramp (ms) | 43 | ( 3 ) | 49 | ( 11 ) | 0.17 | 12.65 | 1.49 | 8.6 |
|  | Laryngeal Complex Ramp (ms) | 55 | ( 7 ) | 74 | ( 12 ) | **0.01** | 3.30 | 3.02 | 11 |
|  | Submental Complex Decay (ms) | 85 | ( 20 ) | 110 | ( 42 ) | 0.26 | 4.28 | 1.19 | 11 |
|  | Laryngeal Complex Decay (ms) | 101 | ( 24 ) | 126 | ( 49 ) | 0.31 | 4.05 | 1.07 | 11 |
|  | Swallow Sequence (ms) | 15 | ( 7 ) | 29 | ( 9 ) | **0.01** | 1.73 | 2.95 | 11 |
|  | SR Diaphragm IBI (ms) | 408 | ( 103 ) | 492 | ( 187 ) | 0.38 | 3.31 | 0.92 | 11 |
|  | SR Inspiratory Delay (ms) | 249 | ( 98 ) | 280 | ( 164 ) | 0.71 | 2.84 | 0.38 | 11 |
|  | Swallow Onset: Insp Peak (ms) | 65 | ( 33 ) | 74 | ( 37 ) | 0.67 | 1.61 | 0.43 | 11 |
|  | Swallow Onset: Insp Onset (0-1 of RC) | 0.24 | ( 0.04 ) | 0.26 | ( 0.06 ) | 0.67 | 1.21 | 0.44 | 11 |

Provides means, standard deviations (SD), *p*-values, F-statistic (F), t-statistics (t), and degrees of freedom (df) from an unpaired t-test comparing A,D) demographics, B,E) swallow characteristics and C,F) swallow-breathing parameters between male and female Vglut2-Ndufs4-KO mice exposed to room air (RA) A-C and chronic hypoxia (CH) D-F. \* Indicates a statistically significance variance (F-test to compare variances), and Welch's t test was used. Swallow related (SR), inter-burst-interval (IBI), grams (g), milliseconds (ms), respiratory cycle (RC).

Furthermore, we examined the impact of CH on the regeneration of respiratory rhythm, following a swallow. A two-way mixed effects ANOVA revealed a significant interaction between genotype and IBI (*p* <0.0001, df = 50, F = 6.66) in RA vs CH Vglut2-Ndufs4-KO mice. Tukey’s multiple comparison test revealed the IBI in Vglut2-Ndufs4-KO RA is significantly longer in duration compared to Vglut2-Ndufs4-KO CH from breaths 1-30 (p< 0.05) (Figure 5A). While there was no statistical difference between the average IBI in control CH and Vglut2-Ndufs4-KO CH mice. We observed a decrease in the variability in Vglut2-Ndufs4-KO CH mice IBI duration compared to Vglut2-Ndufs4-KO RA mice (Figure 5C). Tukey’s multiple comparison test taken from the two-way mixed effects ANOVA (discussed above) revealed no statistical difference in the means or variance in SR IBI and inspiratory delay (breath 1) between control CH mice and Vglut2-Ndufs4-KO CH (Table 3, Figure 5B). These results suggest that exposure to CH prevented abnormal recovery of respiratory rhythm following a swallow in Vglut2-Ndufs4-KO mice but does not fully restore the IBI variability (Figure 5C).

**Figure 5.**
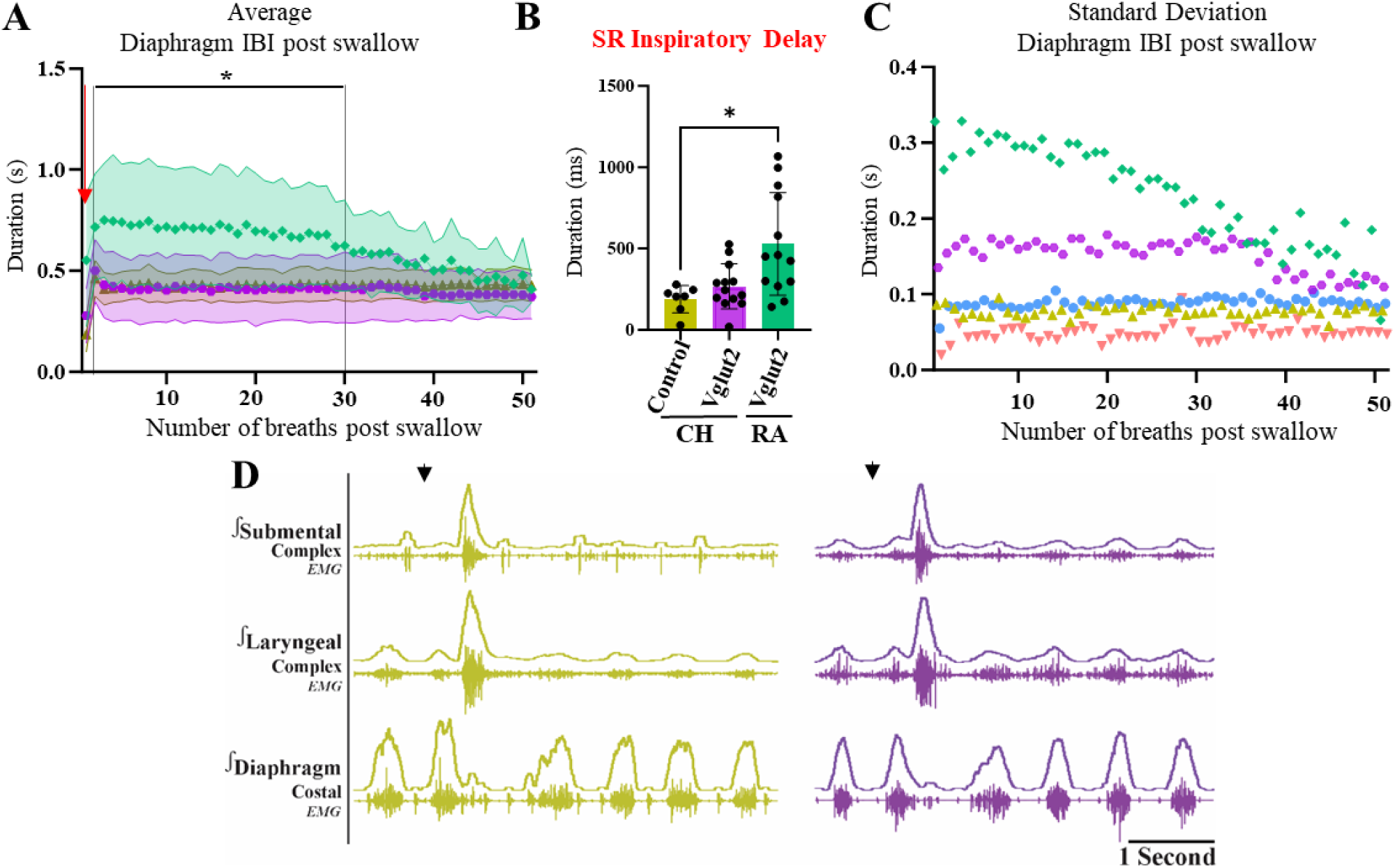
Chronic hypoxia (CH) rescues delayed inspiratory regeneration following a swallow but does not fully restore diaphragm inter-burst-interval (IBI) variability. A) Plot of the average diaphragm IBI following a swallow for control (gold) and Vglut2-Ndufs4-KO (purple) mice exposed to CH and Vglut2-Ndufs4-KO (green) exposed to room air. Vglut2-Ndufs4-KO mice exposed to RA have significantly longer IBI compared to control and Vglut2-Ndufs4-KO mice exposed to CH from breaths. There is no difference in IBI duration between control and Vglut2-Ndufs4-KO mice exposed to CH. The red arrow indicates the first breath post swallow termed B) swallow related (SR) inspiratory delay which is significantly longer in Vglut2-Ndufs4-KO RA mice compared to control CH mice, but there is no difference between Vglut2-Ndufs4-KO and control CH mice. Figure 2 and C) depicts the variability of IBI duration in Vglut2-Ndufs4-KO RA mice (green) which is only partially rescued by CH (purple). We see little differences in the variability between control RA (blue) and CH (gold) mice with the Gad2-Ndufs4-KO RA mice (pink). D) Representative traces of control and Vglut2-Ndufs4-KO mice exposed to CH showing no change in IBI and inspiratory delay in water-triggered swallows.

To distinguish if swallows occur at different times throughout the respiratory cycle, we measured the duration from the peak of the respiration related diaphragm burst, to the start of swallow. Tukey’s multiple comparisons test, taken from a two-way mixed effects ANOVA, revealed swallows in Vglut2-Ndufs4-KO CH mice occur closer to the peak of the diaphragm then Vglut2-Ndufs4-KO RA mice (71 ± 34ms, 273 ± 244ms, *p* = 0.03) (Figure 5-supplemental figure 1A). Though the Vglut2-Ndufs4-KO RA mice have more variability than the other mouse groups. To normalize for respiratory cycle duration, we measured the position of where swallows occur ranging from 0-1. We found swallows occur earlier in the respiratory cycle in Vglut2-Ndufs4-KO CH than control CH mice (0.24 ± 0.08, 0.32 ± 0.04, *p* = 0.01), though there is no difference in timing between the in Vglut2-Ndufs4-KO RA and CH mice (Figure 5-supplemental figure 1B). CH removes the variability of swallow onset in relation to the peak of inspiratory diaphragm activity.

### Sex differences in swallow and breathing characteristics

We found no sex-specific differences in any swallow or swallow-breathing characteristics in control RA and CH mice, Gad2-Ndufs4-KO RA mice, or Vglut2-Ndufs4-KO RA. However, we did see female Vglut2-Ndufs4-KO mice exposed to CH had a longer swallow sequence (29 ± 9ms, 15 ± 7ms, *p* = 0.01) and LC ramp (74 ± 12ms, 55 ± 7ms, *p* = 0.01) than male mice. All data pertaining to sex specific measured parameters are in tables 4-6.

### Presence of microglia in swallow and breathing related medullary regions

Using Iba-1 we stained for microglia in medullary regions related to swallow and breathing such as the nucleus of the Solitary Tract (NTS), ventral respiratory column (VRC) and the postinspiratory complex (PiCo) in control (N=5) and Vglut2cre-Ndufs4-KO (N=6) mice in both RA and CH exposure (Figure 6). A one-way ANOVA with Tukey’s multiple comparison test revealed a significant decrease in the density of Iba-1 positive cells in Vglut2cre-Ndufs4-KO mice in the VRC area compared to RA control mice (0.07 ± 0.009, 0.08 ± 0.008, *p* = 0.05). There was a decrease in the number of Iba-1 positive cells in Vglut2cre-Ndufs4-KO CH mice in PiCo compared to Vglut2cre-Ndufs4-KO RA (54 ± 4, 63 ± 4, *p* = 0.01). Further, we evaluated the morphology of the microglia detected in these regions using morphometric assays, including skeletonization and fractal analysis (Figure 6-figure supplement 1) (29, 30). A one-way ANOVA with Tukey’s multiple comparison test revealed a significant increase in lacunarity of the VRC neurons in Vglut2cre-Ndufs4-KO RA compared to control RA (0.43 ± 0.03, 0.34 ± 0.29, *p* = 0.0004) and Vglut2cre-Ndufs4-KO CH (0.43 ± 0.03, 0.37 ± 0.02, *p* = 0.01). The VRC also had changes in circularity with a significant increase in Vglut2cre-Ndufs4-KO RA compared to control RA (0.88 ± 0.04, 0.78 ± 0.03, *p* = 0.002) and Vglut2cre-Ndufs4-KO CH (0.88 ± 0.04, 0.81 ± 0.03, *p* < 0.0001). We also saw a significant increase in lacunarity of PiCo in Vglut2cre-Ndufs4-KO RA compared to control RA mice (0.42 ± 0.04, 0.35 ± 0.02, *p* = 0.002). However, we did not see any changes in the number or morphology of microglia in the NTS. Taken together this data suggest that chronic hypoxia decreases the number of microglia and restores the lacunarity and circularity of VRC cells.

**Figure 6.**
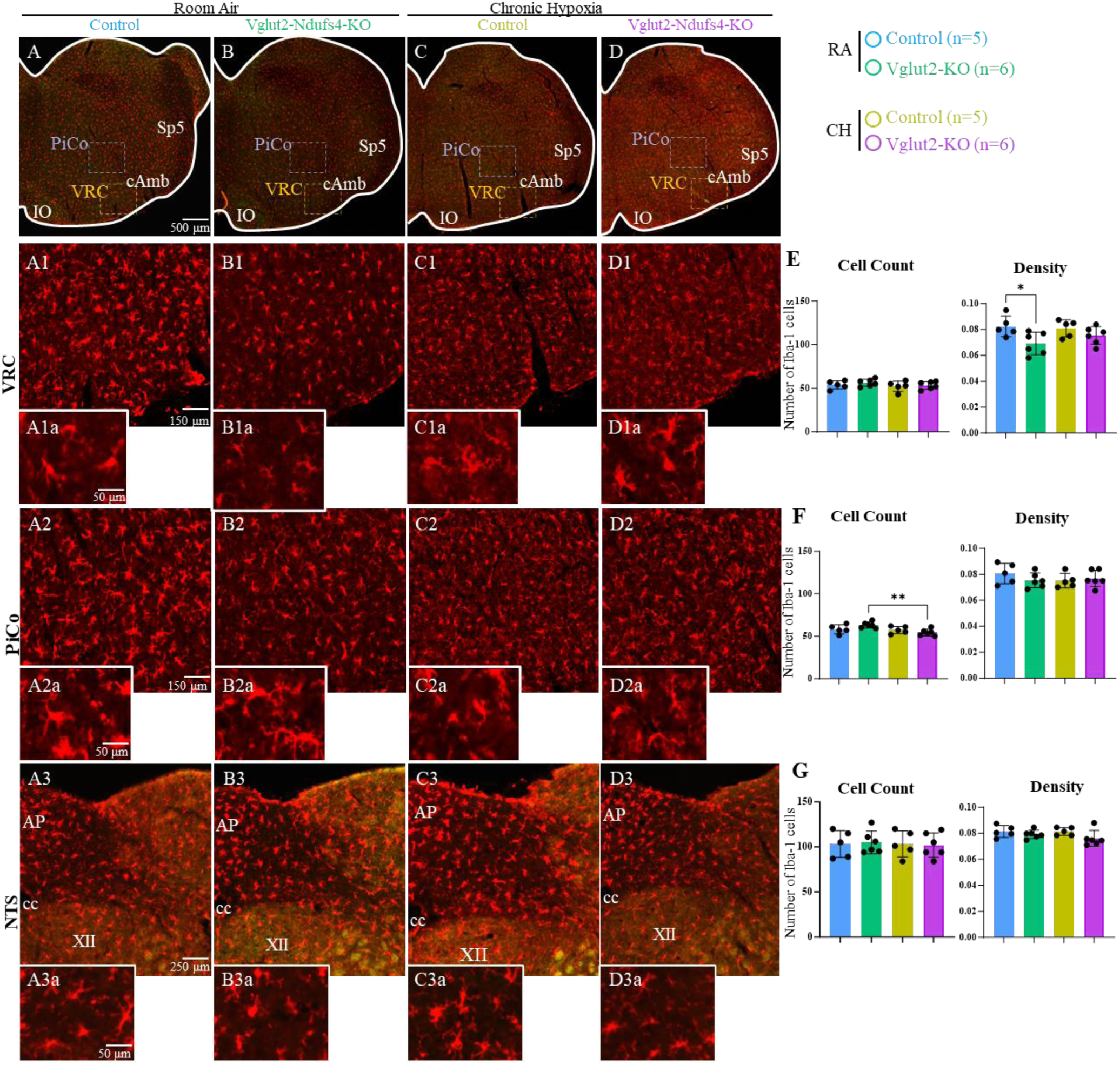
Microglia in breathing and swallow-related medullary centers. Using Iba-1 we stained for microglia in 1) the ventral respiratory column (VRC), 2) postinspiratory complex (PiCo), and 3) the nucleus of the Solitary Tract (NTS) of A) control room air (RA), B) Vglut2-Ndufs4-KO RA, C) control chronic hypoxia (CH), and D) Vglut2-Ndufs4-KO CH mice. A one-way ANOVA with Tukey’s multiple comparison test revealed a significant decrease in E) the density of Iba-1 positive cells in Vglut2cre-Ndufs4-KO mice in the VRC area compared to RA control mice (*p* = 0.05). F) There was a decrease in the number of Iba-1 positive cells in Vglut2cre-Ndufs4-KO CH mice in PiCo compared to Vglut2cre-Ndufs4-KO RA (*p* = 0.01). G) There were no changes to the number or density of microglia in the NTS. Abbreviations: cAmb, nucleus ambiguus pars compacta; Sp5, spinal trigeminal nucleus; AP, area postrema; IO, inferior olive; cc, central canal; XII, hypoglossal nucleus.

## Discussion

Dysphagia and breathing disturbances are typical for children with Leigh syndrome. Management of dysphagia is important in preventing airway obstruction, aspiration, malnutrition, and dehydration leading to weight loss, failure to thrive, aspiration pneumonia, and contribute to spontaneous death (31). Mutations to the NDUFS4 gene cause severe and early-onset LS (18). Here we have described alterations in swallow and breathing activity in Vglut2cre-Ndufs4-KO mice, but not the Gadcre-Ndufs4-KO mice. Homozygous KO of NDUFS4 in glutamatergic neurons resulted in increased variability to the swallow motor pattern, disrupted breathing regeneration, and water-induced apneas. All of which likely contribute to weight loss and premature death seen in this mouse model. Exposure to chronic hypoxia prevented alterations to swallow and breathing physiology which resulted in weight maintenance and life expectancy.

### Variability of swallow motor patterning and regeneration of breathing

Swallowing consists of 3 phases: oral, pharyngeal, and esophageal. It is a complex behavior in which the first two phases share the upper airway with breathing and require precise temporal synchronization from 26 pairs of muscles across 5 cranial nerves to ensure airway protection (32). Alteration of muscle and temporal activity can put a subject at risk of food/liquid entering the airway instead of the esophagus. It is noted that in children with LS that repeated bouts of aspiration pneumonia occur due to feeding and swallowing difficulties which result in gastrostomy and tracheostomy tube placement (13). It is unclear how mutations to the NDUFS4 gene disrupt swallow neuromuscular activity resulting in dysphagia. However, this study has shined light onto the changes in swallow and their interaction with breathing in a mouse model of LS. Activation of the laryngeal complex during swallowing closes the airway to allow food and liquid to safely pass to the esophagus and not the trachea. We saw a significant increase in the amount of time it took the laryngeal complex to fully activate and remain active. However, laryngeal, and submental activation as well as decay and ramp, respectively, were significantly variable across each animal. In healthy mammals, swallow motor pattern must be extraordinarily stable, likely ensured by the efficiency of its command center within the central nervous system. Laryngeal elevation performed by the submental complex and laryngeal closure by the larynx are governed across a vast population of various motor neuron pool, hypoglossal, trigeminal, and facial; nucleus ambiguus. Exposure to chronic intermittent, as opposed to chronic sustained, hypoxia, a method that mimics obstructive sleep apnea or intermittent apneas, creates variability to swallow motor patterning due to changes in the Postinspiratory Complex (PiCo) (24). Suggesting the variability of this motor patterning could originate in several areas. It is likely swallow motor instability which puts the airway at risk and contributes to the weight loss and premature death (33).

Swallow has a hierarchical control over the breathing network. Swallow command areas inhibit ongoing inspiration and when completed allow for regeneration of the respiratory rhythm (22, 23). The automatic and rhythmic nature of breathing allows for efficient oxygen intake and carbon dioxide removal to maintain blood-gas homeostasis. While swallow is a physiological perturbation of the respiratory rhythm, it should not cause prolonged disruption, as seen in this study. Ongoing swallow-breathing discoordination can inhibit continuous feedings and drinking resulting in, prolonged mealtimes, aspiration, and decline in nutrition warranting the need for gastrostomy placement (34). Oral-fed infants should have a normal pharyngoesophageal cardiorespiratory response to swallowing indicated by minimal impact on breathing and heart rate during swallowing. However, we see here in the Vglut2-Ndufs4-KO mice that the breathing rhythm is highly disturbed and requires on average 28 breaths following a swallow to return to normal breathing frequency (35).

### Water-induced apneas disrupt physiological homeostasis in LS

Breath-holding during feeding and swallowing has been shown in Rett Syndrome during spoon feedings (36) and in infants, who have later died of SIDS, during bottle feedings (37, 38). Since these studies focus primarily on nutritive swallowing/sucking, it is unclear whether nonnutritive swallow/sucking, such as sucking on a pacifier or swallowing saliva/mucus for pharyngeal clearance, would also induce apnea. However, regardless of the nutritive behavior, simply touching the larynx or hypopharynx with a laryngoscope will induce apnea to those with increased sensitivity (37). In this study, water administered to the oral cavity, targeting the back of the pharyngeal wall, resulted in apnea with either a swallow (type 1), no upper airway motor activity (type 2), or laryngeal reflex (type 3). Research in various animal models show laryngeal induced apneas occur via laryngeal chemoreflex (LCR), which arises from activation of the superior laryngeal nerve (SLN), and when transected apneas no longer occur (39–41). The physiologic danger resulting from these central apneas are a decrease in O_2_ saturation leading to cyanosis, bradycardia (42), and in immature animal models, more severe apneas resulted in death from asphyxiation (39, 40). Here in this study, we saw on average a 174% decrease in HR when apneas were 20 seconds or longer, whereas in apneas less than 8 seconds only a slight 10% decrease in heart rate occurred, which was not correlated with the duration of apnea. It is possible that this mouse model of LS has some impairment of cardiorespiratory coupling and an abnormal pharyngoesophageal cardiorespiratory response since large changes in HR during apneas of 8s or less did not occur (35, 43). It is also possible that the water-induced apneas seen here could be due to hyperactivity of the SLN. In addition to the unpleasant nature of feeding sessions, dysregulation of cardiorespiratory functions with swallow motor effort leads to an increase in metabolic effort, a decrease in caloric intake and ultimately failure to thrive. These factors result in the necessity of feeding and tracheal tubes, altering the quality of the child’s life (15, 16).

### Chronic hypoxia prevents swallow motor variability and maintains swallow-breathing coordination

Complex I of the mitochondria electron transport chain (ETC) is the first and major entry point for electrons, which oxidizes NADH. It is considered the rate limiting step in cellular respiration and is implicated in the regulation of reactive oxygen species (ROS) (44). Mutation to the NDUFS4 gene leads to dysfunction of complex I in the ETC (20). Mitochondria consume 90% of oxygen in our body (45), however this consumption is declined in those with complex 1 deficiencies such as LS (46). LS mice also have impaired oxygen consumption shown by increased O_2_ levels in the brain. Tissue damage due to oxygen toxicity is likely caused by a mismatch in O_2_ delivery and usage (46). Inspired by the work of Jain et al, who first introduced chronic hypoxia (CH) as a potential therapy for mitochondrial disease in the global KO of NDUFS4 in mice (25), we show for the first time, the effects of CH in mice where NDUFS4 is knocked out specifically in Vglut2 cells on swallow and its coordination with breathing. Similar to Jain et al, Vglut2-Ndufs4-KO mice chronically breathing 11% O_2_ did not experience weight loss or premature death (25). CH prevented variability of the swallow motor pattern and prolonged perturbation of the breathing rhythm. Treatment of global Ndufs4-KO mice with CH (implemented by reducing the delivery of O_2_ to brain tissues, either by 600 ppm of carbon monoxide, 11% oxygen, or anemia) reversed brain lesions, though brain complex I activity remained reduced to similar levels as RA mice and prevented disease progression (25, 26). It was observed that CH does not repair complex I, rather increases complex II activity utilizing fumarate as the electron acceptor instead of oxygen (47), though this remains to be confirmed in Vglut2-Ndufs4-KO mice.

### Cell-specific mitochondrial disruption reveals excitatory neurons play a key role in the production of normal swallow-breathing behavior

There is still much to be discovered about the central control of swallow and its interaction with breathing. Studies have shown the importance of the inspiratory rhythm generator, preBötzinger Complex (preBötC), and the Postinspiratory Complex (PiCo) in the regulation of normal swallow production, laryngeal adduction, and coordination of swallow and breathing (22, 23). Swallow generating neurons are thought to be excitatory by nature, though it has not been determined which brainstem neurons are necessary and sufficient for normal swallow function. Here we show, swallow and breathing disruption in the Vglut2cre-and not the Gad2cre-Ndufs4-KO mice. This suggests the importance of excitatory glutamatergic (Vglut2) neurons in the normal production of swallow and interaction with breathing. It has been shown that Gad2-Ndufs4-KO mice have lesions in the basal ganglia, impaired interneuron excitability in the hippocampus, exhibit severe seizures, mimicking children with LS (28). In contrast, Vglut2-Ndufs4-KO mice have lesions in the brainstem and exhibit breathing and motor impairments (Table 2, (11)). The mechanism behind the region-specific lesions is not understood. Swallow neurons are thought to be under tonic GABAergic inhibition (48). The lack of swallow-breathing disturbance suggests that NDUFS4 mutations in Gad2cre neurons does not disrupt brainstem GABAergic signaling related to control of this function.

### Significance of a pre-clinical mouse model of Leigh Syndrome

The aspiration reflex, colloquially known as a reverse sneeze, is a pharyngeal clearing behavior, generated by a large burst of diaphragm activity creating negative plural pressure. (49, 50) Functionally the forces generated by the diaphragm move mucus or other material from the upper airway to the esophagus and clear the pharynx by triggering a swallow (51, 52). Aspiration of material into the lower airway and lungs should be prevented by laryngeal closure during swallowing and postinspiration. However, methods used here do not allow for observation of the pharyngeal clearing efficacy of AspR in these mice (53). This reflex is experimentally induced by mechanical stimulation to the nasopharynx. However, in this work, aspiration reflex was not purposefully evoked, but triggered in response to a bolus of water in the oral cavity. This is not to be confused with aspiration seen clinically using a Videofluoroscopic swallow study. An aspiration reflex surrounding a swallow does not normally occur in healthy mice or mice exposed to CH, shown in this study. The duration of the diaphragm during the aspiration reflex in the healthy cat was 72 ms with the amplitude reaching 650% of the baseline eupnea activity (49). While we did not see a significant decrease in diaphragm duration and only a maximum of 203% of the diaphragm amplitude, likely due to changes in muscle activity due to the genetic deletion.

### Conclusion

The use of anesthesia eliminates volitional swallowing but allows for detailed analysis of muscle and nerve activity. Videofluoroscopic swallow studies have been adapted in rodents and is necessary to evaluate functional efficacy of swallow in LS mice (54). The combination of anesthetized, present study, and freely behaving swallow studies, creates an opportunity of a preclinical model to test current and future therapeutics for feeding and swallowing disturbances associated with Leigh syndrome. This study introduces the Vglut2-Ndufs4-KO mouse model as a preclinical model to assess swallow and breathing in Leigh Syndrome. Exposure to chronic hypoxia restores swallow and breathing motor patterning, body weight, and life expectancy. Discovery of the mechanisms behind CH-related recovery will be key to finding an effective treatment for Leigh Syndrome.

## Abbreviations

LS: Leigh Syndrome
KO: Knock Out
VRC: Ventral Respiratory Column
PiCo: Postinspiratory Complex
NTS: Nucleus of the Solitary Tract
preBötC: preBötzinger Complex
SC: Submental Complex
LC: Laryngeal Complex
SR: Swallow Related
IBI: inter-burst-interval
HR: Heart Rate
RA: Room Air
CH: Chronic Hypoxia
AspR: Aspiration Reflex
ETC: Electron Transport Chain

## Data Availability

All data is publicly available at 10.6084/m9.figshare.26488018

## Acknowledgements and Funding

We are grateful to receive the NIH grants F32 HL160102 (awarded to A.H.), R01 NS102796 (awarded to F.K.), P01 HL144454 and Project 2 (awarded to J.M.R) HL144801 (awarded to J.M.R.), R01 HL151389 (awarded to J.M.R.), and R01 HL126523 (awarded to J.M.R) for funding this project.

## Author Contributions

A.H., M.K.A., L.M.O., J.M.R. and F.K designed research; A.H., M.K.A., L.M.O., and F.E performed research; A.H., M.K.A., L.M.O., and F.E analyzed data; and A.H., L.M.O., J.M.R., and F.K wrote the paper.

---

**Figure 1-Supplemental Figure 1.**
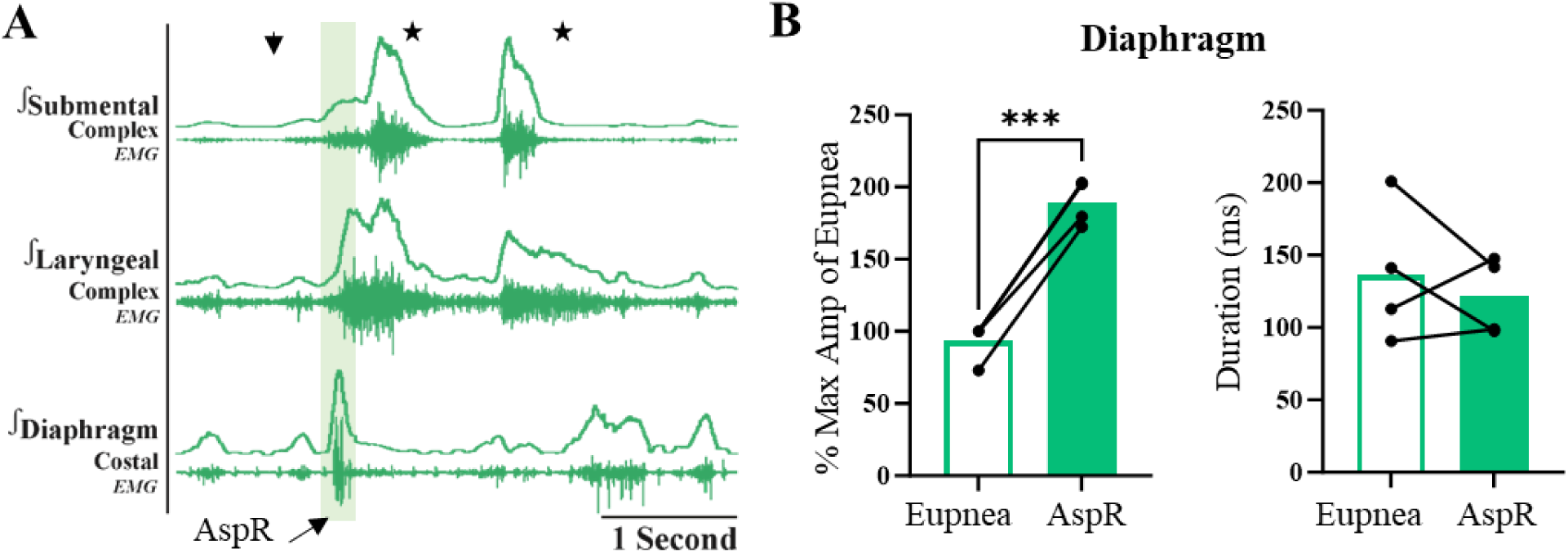
Airway protective behavior, likely an aspiration reflex (AspR), in response to water in Vglut2-Ndufs4-KO mice. A) Representative trace of water (down arrow) induced swallow (stars) and AspR indicated in the diaphragm. This airway protective behavior is seen only in the LS mouse model with disordered swallow, the Vglut2-Ndufs4-KO mice exposed to room air. AspR is triggered by stimulation of the pharyngeal mucosa described as a strong and rapid diaphragm activation presumably strong enough to remove material from the oropharynx and clear the airway via reflexive swallowing (53). B) There was a significant increase in diaphragm amplitude normalized to the maximum of the previous eupnea diaphragm burst (182 ± 12 vs 90 ± 13m % max of eupnea, *p* = 0.0001). Though we saw no change in diaphragm duration, this could be attributed to the disease.

**Figure 5-Supplemental Figure 1.**
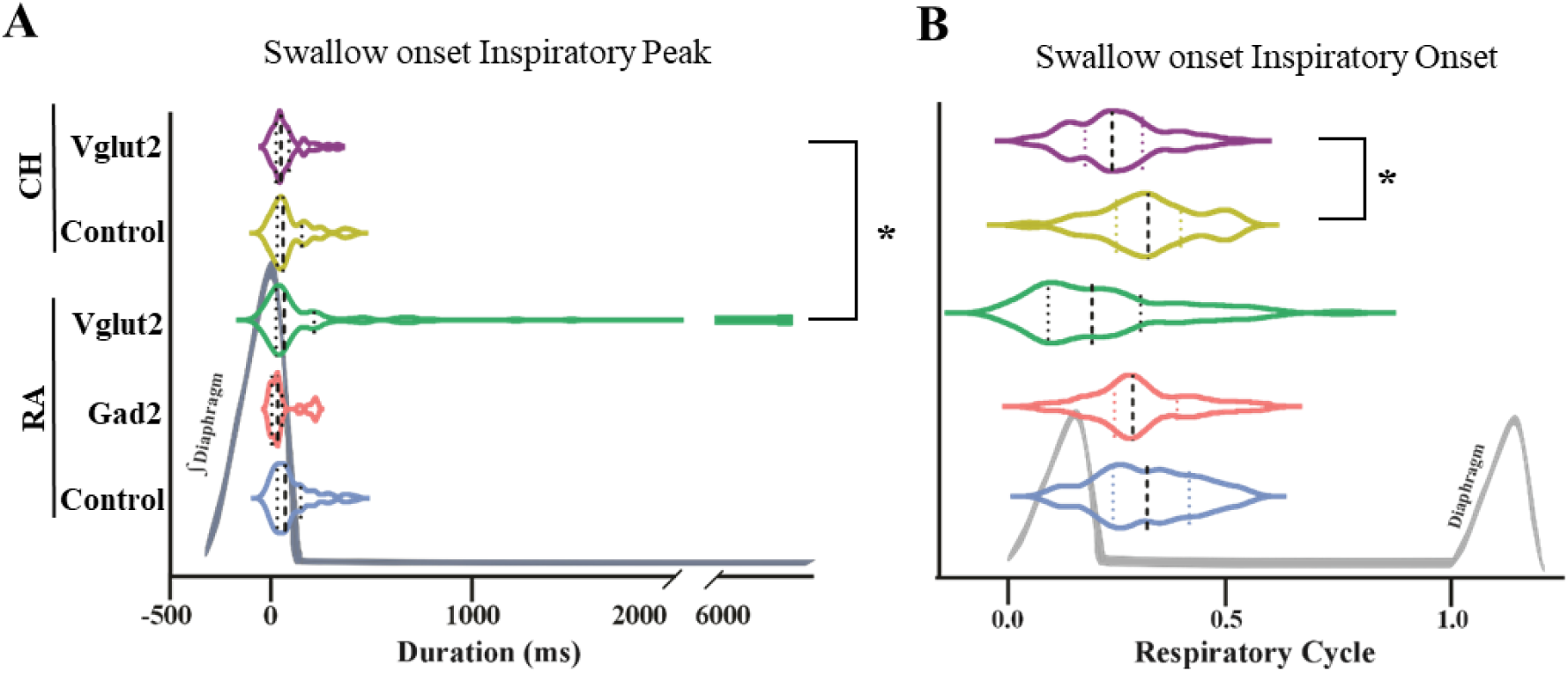
Swallow timing in relation to breathing in all genetic mouse types exposed to room air (RA) and chronic hypoxia (CH). We saw a significant difference in swallows occurring after the peak of the diaphragm and greater variability in Vglut2-Ndufs4-KO mice exposed to RA compared to CH (*p =* 0.03). B) when normalized to the respiratory cycle, there was a significant difference with swallows occurring sooner in the respiratory cycle for Vglut2-Ndufs4-KO CH mice compared to control CH mice (*p* = 0.01) (Table 3).

**Figure 6-Figure Supplement 1.**
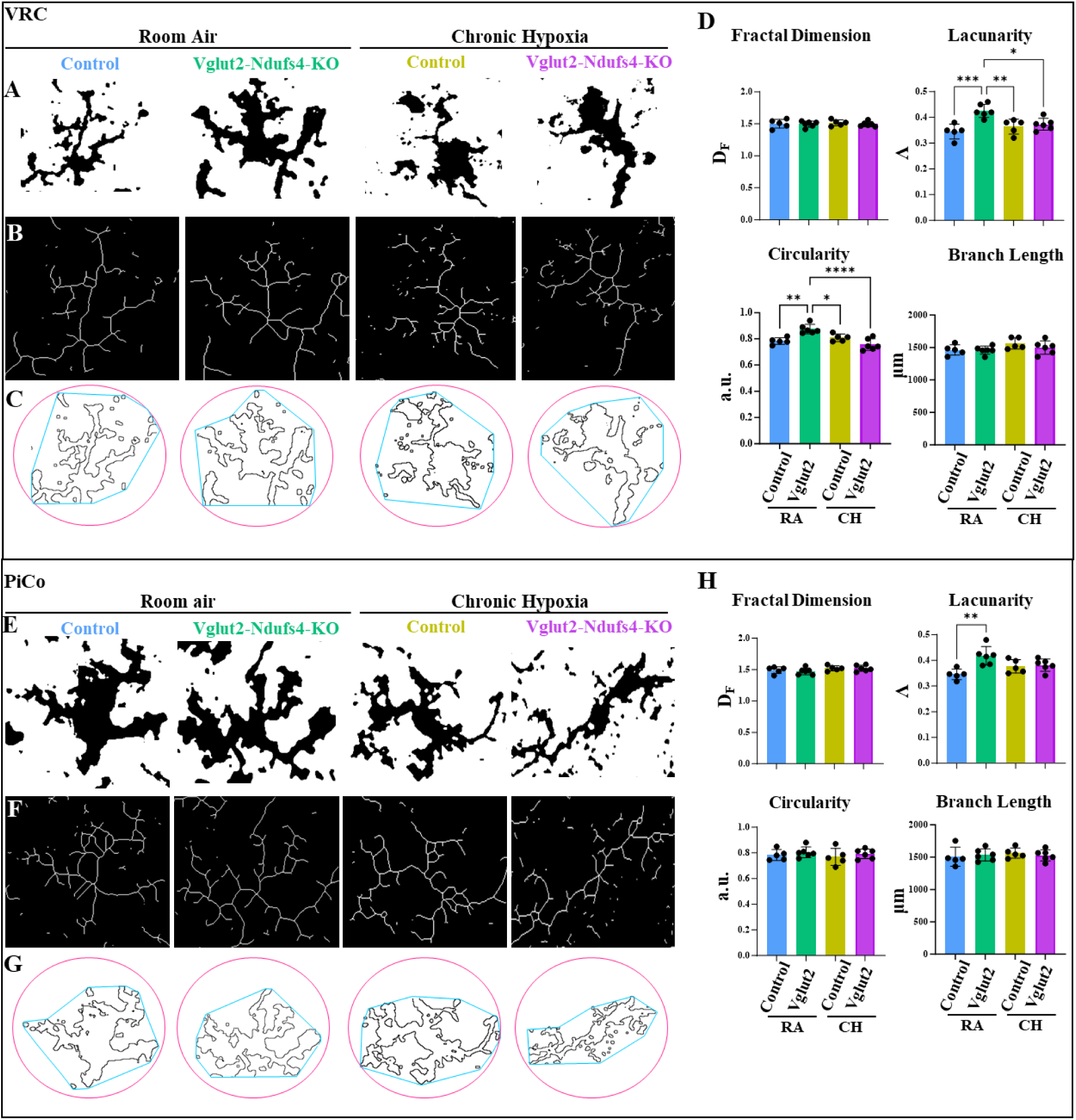
Morphologic changes to microglia in breathing and swallow-related medullary centers using morphometric assays. A) Black and white contrast, B) skeletonization, and C) circularity of a microglia in each genetic and oxygen condition in the ventral respiratory column (VRC). D) A one-way ANOVA with Tukey’s multiple comparison test revealed a significant increase in lacunarity of the VRC neurons in Vglut2cre-Ndufs4-KO RA compared to control RA (*p* = 0.0004) and Vglut2cre-Ndufs4-KO CH (*p* = 0.01). The VRC also had changes in circularity with a significant increase in Vglut2cre-Ndufs4-KO RA compared to control RA (*p* = 0.002) and Vglut2cre-Ndufs4-KO CH (*p* < 0.0001). E) Black and white contrast, F) skeletonization, and G) circularity of a microglia in each genetic and oxygen condition in the postinspiratory complex (PiCo). H) There was significant increase in lacunarity of PiCo in Vglut2cre-Ndufs4-KO RA compared to control RA mice (*p* = 0.002).

## References

1. Safi, F., et al., P653 Swallowing disorder revealing leigh’s syndrome. 2019, BMJ Publishing Group Ltd.

2. Montpetit, V.J., et al., Subacute necrotizing encephalomyelopathy: a review and a study of two families. Brain, 1971. 94(1): p. 1–30.

3. Leigh, D., Subacute necrotizing encephalomyelopathy in an infant. Journal of neurology, neurosurgery, and psychiatry, 1951. 14(3): p. 216.

4. Rahman, S., et al., Leigh syndrome: clinical features and biochemical and DNA abnormalities. Annals of Neurology: Official Journal of the American Neurological Association and the Child Neurology Society, 1996. 39(3): p. 343–351.

5. Hong, C.-M., et al., Clinical characteristics of early-onset and late-onset Leigh syndrome. Frontiers in neurology, 2020. 11: p. 267.

6. Chuquilin, M., et al., Response to immunotherapy in a patient with adult onset Leigh syndrome and T9176C mtDNA mutation. Molecular Genetics and Metabolism Reports, 2016. 8: p. 28–32.

7. Piao, Y.S., et al., Clinico-neuropathological study of a Chinese case of familial adult Leigh syndrome. Neuropathology, 2006. 26(3): p. 218–221.

8. Hayashi, Y., et al., Clinicopathological findings of a mitochondrial encephalopathy, lactic acidosis, and stroke-like episodes/Leigh syndrome overlap patient with a novel m. 3482A> G mutation in MT-ND1. Neuropathology, 2021. 41(1): p. 84–90.

9. Sofou, K., et al., A multicenter study on Leigh syndrome: disease course and predictors of survival. Orphanet journal of rare diseases, 2014. 9(1): p. 1–16.

10. Kistol, D., et al., Leigh Syndrome: Spectrum of Molecular Defects and Clinical Features in Russia. International Journal of Molecular Sciences, 2023. 24(2): p. 1597.

11. Bolea, I., et al., Defined neuronal populations drive fatal phenotype in a mouse model of Leigh syndrome. Elife, 2019. 8: p. e47163.

12. Stenton, S.L., et al., Leigh syndrome: a study of 209 patients at the Beijing Children’s hospital. Annals of Neurology, 2022. 91(4): p. 466–482.

13. Tanaka, R., et al., Novel NARS2 variant causing leigh syndrome with normal lactate levels. Human Genome Variation, 2022. 9(1): p. 12.

14. Martin, E., et al., Brainstem lesion revealed by MRI in a case of Leigh’s disease with respiratory failure. Pediatric Radiology, 1990. 20: p. 349–350.

15. Gozal, D., et al., Leigh syndrome: anesthetic management in complicated endoscopic procedures. Pediatric Anesthesia, 2006. 16(1): p. 38–42.

16. Yashon, D. and J.A. Jane, Subacute necrotizing encephalomyelopathy of infancy and childhood. Journal of Clinical Pathology, 1967. 20(1): p. 28.

17. Ramelli, G.P., et al., Gastrostomy placement in paediatric patients with neuromuscular disorders: indications and outcome. Developmental Medicine & Child Neurology, 2007. 49(5): p. 367–371.

18. Ortigoza-Escobar, J.D., et al., Ndufs4 related Leigh syndrome: A case report and review of the literature. Mitochondrion, 2016. 28: p. 73–78.

19. Calvaruso, M.A., et al., Mitochondrial complex III stabilizes complex I in the absence of NDUFS4 to provide partial activity. Human molecular genetics, 2012. 21(1): p. 115–120.

20. Kruse, S.E., et al., Mice with Mitochondrial Complex I Deficiency Develop a Fatal Encephalomyopathy. Cell Metabolism, 2008. 7(4): p. 312–320.

21. Martin-Harris, B., et al., Breathing and swallowing dynamics across the adult lifespan. Arch Otolaryngol Head Neck Surg, 2005. 131(9): p. 762–70.

22. Huff, A., et al., Optogenetic stimulation of pre–Bötzinger complex reveals novel circuit interactions in swallowing–breathing coordination. Proceedings of the National Academy of Sciences, 2022. 119(29): p. e2121095119.

23. Huff, A.D., et al., Role of the postinspiratory complex in regulating swallow-breathing coordination and other laryngeal behaviors. Elife, 2023. 12: p. e86103.

24. Huff, A.D., et al., Chronic Intermittent Hypoxia reveals role of the Postinspiratory Complex in the mediation of normal swallow production. Elife, 2024. 12: p. RP92175.

25. Jain, I.H., et al., Hypoxia as a therapy for mitochondrial disease. Science, 2016. 352(6281): p. 54–61.

26. Ferrari, M., et al., Hypoxia treatment reverses neurodegenerative disease in a mouse model of Leigh syndrome. Proceedings of the National Academy of Sciences, 2017. 114(21): p. E4241–E4250.

27. Basmajian, J.V. and G. Stecko, A new bipolar electrode for electromyography. Journal of Applied Physiology, 1962. 17(5): p. 849–849.

28. Manning, A., et al., Elevated susceptibility to exogenous seizure triggers and impaired interneuron excitability in a mouse model of Leigh syndrome epilepsy. Neurobiology of disease, 2023. 187: p. 106288.

29. Morrison, H., et al., Quantitative microglia analyses reveal diverse morphologic responses in the rat cortex after diffuse brain injury. Sci Rep, 2017. 7(1): p. 13211.

30. SheikhBahaei, S., et al., Morphometric analysis of astrocytes in brainstem respiratory regions. J Comp Neurol, 2018.

31. Murry, T., R.L. Carrau, and K. Chan, Clinical management of swallowing disorders. 2020: Plural Publishing.

32. Doty, R.W. and J.F. Bosma, An electromyographic analysis of reflex deglutition. J Neurophysiol, 1956. 19(1): p. 44–60.

33. Feroah, T.R., et al., Effects of spontaneous swallows on breathing in awake goats. Journal of Applied Physiology, 2002. 92(5): p. 1923–1935.

34. van den Engel-Hoek, L., et al., Feeding and swallowing disorders in pediatric neuromuscular diseases: an overview. Journal of neuromuscular diseases, 2015. 2(4): p. 357–369.

35. Hasenstab, K.A. and S.R. Jadcherla, Evidence-based approaches to successful oral feeding in infants with feeding difficulties. Clinics in Perinatology, 2022. 49(2): p. 503–520.

36. Morton, R.E., et al., Respiration patterns during feeding in Rett syndrome. Developmental Medicine & Child Neurology, 1997. 39(9): p. 607–613.

37. Steinschneider Jr, A. and D.D. Rabuzzi Jr, Apnea and airway obstruction during feeding and sleep. The Laryngoscope, 1976. 86(9): p. 1359–1366.

38. Steinschneider, A., S.L. Weinstein, and E. Diamond, The sudden infant death syndrome and apnea/obstruction during neonatal sleep and feeding. Pediatrics, 1982. 70(6): p. 858–863.

39. Sutton, D., E.M. Taylor, and R.C. Lindeman, Prolonged apnea in infant monkeys resulting from stimulation of superior laryngeal nerve. Pediatrics, 1978. 61(4): p. 519–27.

40. Downing, S.E. and J.C. Lee, Laryngeal chemosensitivity: a possible mechanism for sudden infant death. Pediatrics, 1975. 55(5): p. 640–649.

41. Xia, L., J.C. Leiter, and D. Bartlett Jr, Laryngeal apnea in rat pups: effects of age and body temperature. Journal of Applied Physiology, 2008. 104(1): p. 269–274.

42. Guilleminault, C. and S. Coons, Apnea and bradycardia during feeding in infants weighing greater than 2000 gm. J Pediatr, 1984. 104(6): p. 932–5.

43. Dick, T.E., et al., Cardiorespiratory coupling: common rhythms in cardiac, sympathetic, and respiratory activities. Prog Brain Res, 2014. 209: p. 191–205.

44. Sharma, L.K., J. Lu, and Y. Bai, Mitochondrial respiratory complex I: structure, function and implication in human diseases. Curr Med Chem, 2009. 16(10): p. 1266–77.

45. Rich, P., Chemiosmotic coupling: the cost of living. Nature, 2003. 421(6923): p. 583–583.

46. Jain, I.H., et al., Leigh syndrome mouse model can be rescued by interventions that normalize brain hyperoxia, but not HIF activation. Cell metabolism, 2019. 30(4): p. 824–832. e3.

47. Spinelli, J.B., et al., Fumarate is a terminal electron acceptor in the mammalian electron transport chain. Science, 2021. 374(6572): p. 1227–1237.

48. Wang, Y. and D. Bieger, *Role of solitarial GABAergic mechanisms in control of swallowing.* American Journal of Physiology-Regulatory, Integrative and Comparative Physiology, 1991. 261(3): p. R639–R646.

49. Poliacek, I., et al., Activity of the laryngeal abductor and adductor muscles during cough, expiration and aspiration reflexes in cats. Physiological research, 2003. 52(6): p. 749–762.

50. Korpáš, J. and Z. Tomori, Cough and Other Respiratory Reflexes./Kašeľ a Iné Respiračné Reflexy. 1979: Veda.

51. Tomori, Z., Pleural, tracheal and abdominal pressure variations in defensive and pathologic reflexes of the respiratory tract. Physiologia bohemoslovenica, 1965. 14: p. 84–95.

52. Tomori, Z. and J. Widdicombe, Muscular, bronchomotor and cardiovascular reflexes elicited by mechanical stimulation of the respiratory tract. The Journal of physiology, 1969. 200(1): p. 25–49.

53. Tomori, Z., R. Benacka, and V. Donic, Mechanisms and clinicophysiological implications of the sniff- and gasp-like aspiration reflex. Respir Physiol, 1998. 114(1): p. 83–98.

54. Lever, T.E., et al., Adapting human videofluoroscopic swallow study methods to detect and characterize dysphagia in murine disease models. JoVE (Journal of Visualized Experiments), 2015(97): p. e52319.

55. Pitts, T., et al., Neurons in the dorsomedial medulla contribute to swallow pattern generation: Evidence of inspiratory activity during swallow. PloS one, 2018. 13(7): p. e0199903.

56. Pitts, T. and K.E. Iceman, Deglutition and the regulation of the swallow motor pattern. Physiology, 2023. 38(1): p. 10–24.

